# Insight into differing decision-making strategies that underlie cognitively effort-based decision making using computational modeling in rats

**DOI:** 10.1101/2023.09.20.558692

**Authors:** Claire A. Hales, Mason M. Silveira, Lucas Calderhead, Leili Mortazavi, Brett A. Hathaway, Catharine A. Winstanley

**Author notes:** Correspondence should be addressed to Dr. Claire A. Hales or Dr. Catharine A. Winstanley, Djavad Mowafaghian Centre for Brain Health, Department of Psychology, 2215 Wesbrook Mall, Vancouver, BC, V6T 1Z3, Canada, or. Current address: Fashion Business School, London College of Fashion, University of the Arts London, London, UK. Current address: Department of Psychology, Stanford University, Stanford, CA, USA.

## Abstract

**Rationale:** The rat Cognitive Effort Task (rCET), a rodent model of cognitive rather than physical effort, requires animals to choose between an easy or hard visuospatial discrimination, with a correct hard choice more highly rewarded. Like in humans, there is stable individual variation in choice behavior. In previous reports, animals were divided into two groups - workers and slackers - based on their mean preference for the harder option. Although these groups differed in their response to pharmacological challenges, the rationale for using this criterion for grouping was not robust.

**Methods:** We collated experimental data from multiple cohorts of male and female rats performing the rCET, and used a model-based framework combining drift diffusion modeling with cluster analysis to identify the decision-making processes underlying variation in choice behavior.

**Results:** We verified that workers and slackers are statistically different groups, but also found distinct intra-group profiles. These subgroups exhibited dissociable performance during the attentional phase, linked to distinct decision-making profiles during choice. Reanalysis of previous pharmacology data using this model-based framework showed that serotonergic drug effects were explained by changes in decision boundaries and non-decision times, whilst scopolamine’s effects were driven by changes in decision starting points and rates of evidence accumulation.

**Conclusions:** Modeling revealed the decision-making processes that are associated with cognitive effort costs, and how these differ across individuals. Reanalysis of drug data provided insight into the mechanisms through which different neurotransmitter systems impact cognitively effortful attention and decision-making processes, with relevance to multiple psychiatric disorders.

## Introduction

Decision making often involves determining whether the desired outcome is worth the cognitive effort required to attain it. In our everyday lives, the amount of effort that a person is willing to allocate influences how successful pursuit of a goal is likely to be. An individual must balance assignment of finite mental resources to decisions, tasks and goals. Alterations in cost/benefit decision making have been observed in people living in poverty (Haushofer & Fehr, 2014), and motivational deficits and inappropriate use of attentional resources during decision making are common features in most psychiatric illnesses (Gleichgerrcht et al., 2010; Goschke, 2014; Der-Avakian et al., 2016). Cognitive effort is non-physical and intrinsically costly, incorporating psychological constructs such as attention, response inhibition, cognitive flexibility and working memory (Kool et al., 2010). Many different tasks and paradigms are used in humans to investigate the neural mechanisms and circuitry underlying cognitive effort (e.g. Kool et al., 2010; Dixon & Christoff, 2012; Westbrook et al., 2013; Reddy et al., 2015; Chong et al., 2017), whilst in rodents, the rodent Cognitive Effort Task (rCET) has been used to directly examine how specific brain areas, neurotransmitters and neuromodulators regulate decisions requiring cognitive effort (Cocker et al., 2012; Hosking et al., 2014; 2015; 2016; Silveira et al., 2017; 2018; 2020; 2021).

The rCET is a cognitively demanding attentional task in which rats choose between an easy or hard visuospatial discrimination (Cocker et al., 2012). The task is based on the well-validated five-choice serial reaction-time task (5CSRT) that measures visuospatial attention and motor impulsivity (Robbins 2002), but also includes a choice element for allocation of cognitive effort. On each trial, rats can choose either a trial that requires successful completion of an easier attentional challenge (localisation of a 1 s stimulus light) for lesser reward, or a harder attentional challenge (localisation of a 0.2 s stimulus light) for greater reward (Cocker et el., 2012). Similar to humans (McGuire & Botvinick, 2010), some rats are willing to expend more cognitive effort to earn the larger reward on the rCET, choosing the harder, high effort, high reward (HR) option more frequently. Using a median split, these have been designated “workers”, whilst “slackers” are those rats which choose the low effort, low reward (LR) option relatively more often (Cocker et el., 2012). Although workers and slackers differ in how much cognitive effort they are willing to expend, previous analyses of single cohorts have found they do not differ in other behavioral measures, such as accuracy to detect the stimulus or responses made prematurely before the stimulus light is illuminated, suggesting cognitive effort allocation is not driven by differences in attentional performance or motor impulsivity (Cocker et el., 2012). Pharmacological studies have found that some drugs, including amphetamine, caffeine and nicotine (Cocker et al., 2012) and scopolamine (Hosking et al., 2014, replicated by Silveira, 2018) differentially alter HR choice in workers versus slackers, whilst others act similarly on both groups (e.g. serotonergic drugs; Silveira et al., 2020). It has been proposed that slackers might be more sensitive to cognitive effort costs, experiencing a greater sense of effort on HR trials than workers (Cocker et al., 2012). Use of different strategies to perform the task may also contribute, allowing workers to display equivalent levels of accuracy to slackers, whilst still overcoming the higher cognitive effort demands (Cocker et al., 2012). From traditional analyses of standard behavioral measures, it is not possible to probe these possibilities further.

One way to investigate this is with cognitive models. Cognitive models simulate the mental processes that might underlie a particular behaviour, such as decision making, and provide a framework for predicting and comprehending performance on a task in terms of the underlying cognitive processes. The drift diffusion model (DDM; Ratcliff, 1978), a model of two-choice decision making, has been widely used to investigate the processes underlying perceptual decisions (see Voss et al., 2013 for a review), and more recently successfully applied to preferential or value-based decisions (Dutilh & Rieskamp, 2016; Tajima et al., 2016). The key tenet of the DDM is that a decision is made through accumulation of information over time until a threshold is reached, and the model maps the cognitive processes that would contribute to this type of mechanism onto psychologically meaningful parameters (Ratcliff, 1978). These parameters include: the decision starting point, which represents a predetermined bias for one or other decision; the drift rate, which denotes the rate or strength of information accumulation; boundary separation, which signifies the distance between the two thresholds that each correspond to a choice, and the non-decision time, which incorporates the amount of the total response time (RT) that is due to processes not directly related to the decision, such as sensory and motor processing, and other extraneous factors (Ratcliff & Tuerlinckx, 2002). For value-based decisions, such as the choice between the HR and LR option in the rCET, it is assumed that the information being accumulated is internally generated in the brain based on, for example, prior experience, previously learnt task features, current preferences and physiological state. The drift rate therefore represents how strongly the organism weighs and integrates this internal information, rather than how closely information coming in from the environment matches the organism’s representation of that stimuli, as would be the case in a perceptual task (Milosavljevic et al., 2010).

Utilizing data from multiple cohorts of male and female rats trained on the rCET, the aim of this this work was to fit the DDM to behavioral data, and combine this modeling approach with cluster analysis to: a) determine what decision making processes contribute to choices based on evaluation of cognitive effort; b) investigate whether worker and slacker groupings classified using a median split approach is a valid way to group the data; c) ask are there sex differences in the way that males and females perform the task; and d) re-analyze two previously published pharmacological drug studies to provide further insight into the interaction between neurotransmitter systems and cognitive mechanisms involved in effort-based decision making. We used data from drugs targeting serotonin (5-HT; Silveira et al., 2020) and acetylcholine (Silveira; 2018), receptors, as both neurotransmitter systems have been broadly implicated in decision making (Homberg, 2012; Fobbs & Mizumori, 2014), but with effects on different aspects of this process.

## Methods

### Task

The rCET is a visuospatial discrimination task based on the 5CSRT. On each trial, rats select a lever corresponding to an easy or hard attentional challenge. Following selection of the easy, low reward option (LR) rats have to correctly nosepoke in one of the five stimulus holes to receive a single sugar pellet reward, the correct hole being signaled by a 1 s stimulus light. Conversely, on hard trials (HR), rats can earn two sugar pellets for a correct nosepoke, but the stimulus presentation is much shorter (0.2 s). Although more difficult, hard trials are more advantageous as rats can earn double the reward for successful completion. Within a population of rats, there is much individual variation in the proportion of LR vs HR trials chosen in a session, presumably corresponding to different levels of cognitive effort individuals are willing to allocate to completing the task. Previous work has used a cut-off of 70% to label rats as “workers”, those who choose HR > 70% of the time, and “slackers”, those who choose HR on ≤ 70% of trials, based on the population average being around this value (Cocker et al., 2012; Hosking et al., 2014; 2015; 2016; Silveira et al., 2017; 2018; 2020; 2021). Accuracy of detection of the shorter HR stimulus, as well as how often they fail to wait for stimuli to appear (premature responding) are not different between workers and slackers in individual cohorts, suggesting that cognitive effort allocation is not dependent on task performance. Figure 1 depicts the behavioral measures that are recorded and output in the rCET, and which of these are used for modeling.

**Fig. 1-.**
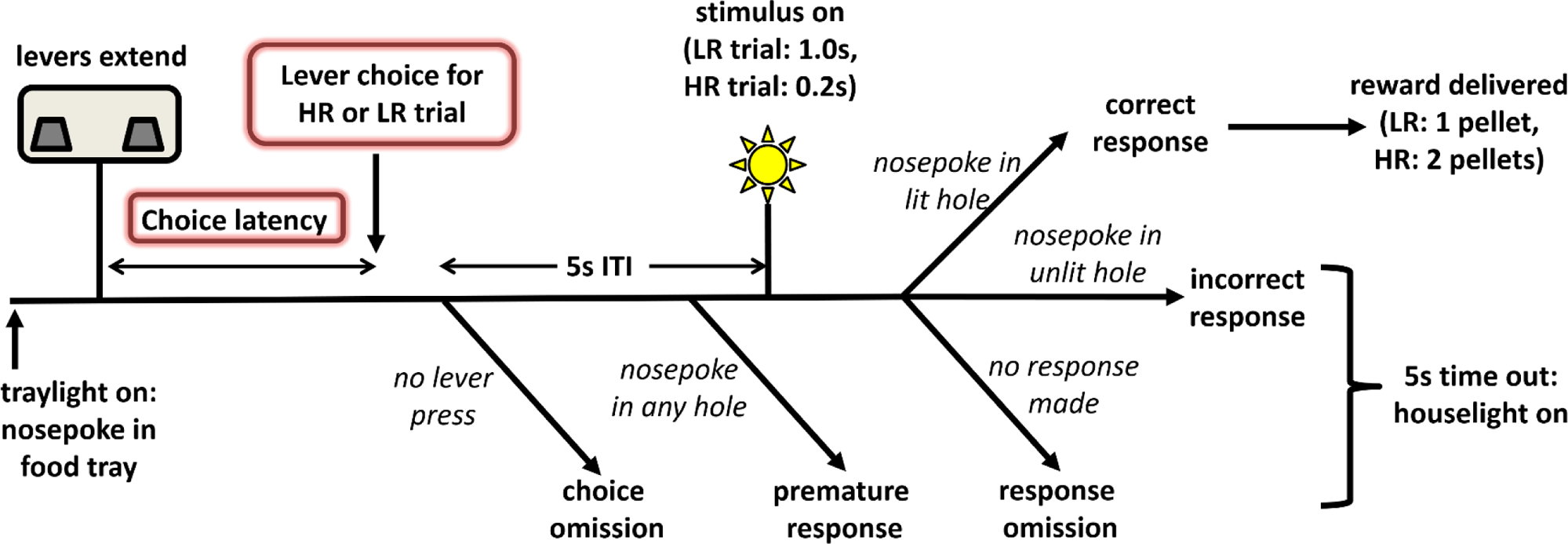
Task schematic depicting the rodent Cognitive Effort Task. The trial format of the rCET is depicted. Briefly, rats initiate a trial with a nosepoke in the food tray. This causes the levers to extend, allowing the rat to choose either a high effort, high reward (HR) or low effort, low reward (LR) trial. No choice results in a choice omission. Following a choice, there is a 5 second intertrial interval (ITI), during which the rat has to wait before the stimulus light comes on. Failure to wait is classed as a premature response. A correct response is made by a nosepoke in the lit hole, following which the appropriate reward is delivered. Incorrect responses are those made in an unlit hole, with an omission occurring if there is no response. Both types of omissions, prematures and incorrect responses are punished with a 5 second time out. Lever choice and choice latency (highlighted with a red box) are the data input into the diffusion model.

### Subjects and Data

Behavioral data used was from six cohorts of rats trained on the rCET (n = 162), three cohorts of male Long Evans rats (total n = 80) and three cohorts of female Long Evans rats (total n = 82). Baseline data from the final three stable acquisition sessions was used from all cohorts for all model and cluster analyses. Data from one female cohort (n = 24; Experiment 1) comprising serotonergic drug manipulations previously published by Silveira et al. (2020), and from one male cohort (n = 39; Experiment 2) comprising an unpublished replication of a scopolamine drug study (original study published by Hosking et al., 2014) was reanalyzed using the DDM and cluster model framework to further probe how these drugs alter decision-making processes. Drugs used in Experiment 1 were: Ro-60-0175, a selective 5-HT_2C_ receptor agonist (0.0, 0.1, 0.3 and 0.6 mg/kg); M100907, a 5-HT_2A_ receptor antagonist (0.0, 0.01, 0.03 and 0.1 mg/kg); SB-242084, a selective 5-HT_2C_ receptor antagonist (0.0, 0.1, 0.25 and 0.5 mg/kg) and 8-OH-DPAT, a 5-HT_1A_ receptor agonist (0.0, 0.1, 0.3 and 0.6 mg/kg). In experiment 2, the mAChR antagonist scopolamine was given at 0.0, 0.03, 0.1, 0.3 mg/kg. Each drug manipulation was carried out as a Latin Square design with the four drug doses given over a two-week period, and were preceded and followed by a week of baseline behavioral testing with no drug administration. Behavioral measures from baseline sessions are also presented. These behavioral measures were: total choices, number of choice omissions, choice latencies for HR and LR trials, percentage of premature responses made after a lever choice, and accuracy to detect the stimulus on HR or LR trials.

### Modeling

The diffusion model was fit to behavioral data from the rCET using fast-dm-30.2 (Voss & Voss, 2007; 2008; Voss et al., 2010; 2015). Lever choice, corresponding to a choice for either a HR or LR trial, and RTs for these choices were input to the model. Figure 1 shows the task structure, and which behavioral measures from the task were used for modeling. Fast-dm calculates predictive cumulative distribution functions (CDFs) for choices and RTs, and then uses a partial differentiation equation solver to model the evolution of the probability distribution forward in time. Parameters are optimised by using an implementation of the Nelder-Mead method (Nelder & Mead, 1965). Further details about diffusion modelling using fast-dm can be found in Voss et al. (2015). Multiple models were tested using different combinations of parameters that were fit to all trials during which a lever choice for the HR or LR option was made to identify the parameter combination that produced best model fits. Validation of this best fitting model was carried on behavioral data from the final three stable acquisition sessions for rats in Experiments 1 and 2 (n = 63). This was then confirmed to also be good fit for behavioural data from other cohorts. Data from individual rats were modeled separately. As carried out previously (Hales et al., 2016; 2017), and following recommendations given in Voss et al. (2015), model fit was assessed using Kolmogorov-Smirnov (KS) test statistics output by fast-dm-30.2. The KS test statistic is the maximum absolute vertical distance between the empirical and the predicted CDFs of the RT distributions. For multiple trials in a task, n, it is computed as:

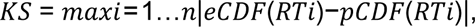

where eCDF and pCDF are the empirical and predicted CDFs, respectively, and RTi is the response latency in trial i. p-values < 0.05 indicate that the model does not demonstrate a good fit. Subjects with model fit KS statistics p < 0.05 were excluded from analyses. Table 1 details the number of animals removed from each cohort, and the reasons for exclusion. The parameter combination that produced the best model fit was selected and used to model the behavioral data. This best fitting model had five parameters: starting point (zr), boundary separation (a), drift rate (v), non-decision RT (t0), and variability in the starting point (szr). The other parameters that can be fit within fast-dm-30.2: the difference in speed of response execution between the two responses (d), variability in drift rate (sv), variability in non-decision RT (st0) and percentage of contaminants (p); were set to 0. In the model, the upper boundary represents a decision for a HR trial, while the lower boundary represents a decision for a LR trial.

**Table 1-.** Numbers of rats for rCET behavioral data used for modeling.

| COHORT | N | SEX | # WORKERS | # SLACKERS | EXCLUDED | FINAL N |
| --- | --- | --- | --- | --- | --- | --- |
| **1** | 11 | M | 6 | 5 | 1 (worker) | 10 |
| **2** | 38 | M | 19 | 19 | 1 (worker) | 37 |
| **3** | 31 | M | 16 | 15 | 2 (workers) | 29 |
| **M TOTAL** | **80** |  | **37 (41)** | **39** | **4** | **76** |
| **4** | 28 | F | 14 | 14 | 0 | 28 |
| **5** | 30 | F | 17 | 13 | 0 | 30 |
| **6** | 24 | F | 11 | 13 | 1 (worker) | 23 |
| **F TOTAL** | **82** |  | **41 (42)** | **40** | **1** | **81** |
| **TOTAL** | **162** |  | **78 (83)** | **79** | **5** | **157** |

Numbers in bold denote summary totals for group or sex. Numbers in brackets are total before exclusions.

For main analyses all five parameters (zr, a, v, t0 and szr) were fit to all trials within a session (all trials condition). All rats excluded (n = 9) due to poor model fits were workers (Table 1). Visual inspection of data from these animals indicated they were nearly always (often around 95%, but for some sessions up to 100% of the time) choosing the HR option, meaning there were insufficient data points in the LR RT distribution for a good model fit (data not shown).

For outcome analyses, all parameters except szr were fit to trials split by whether the rat proceeded to make either a premature response, correctly detect the stimulus, or nosepoke in the incorrect hole (outcome condition). Trials for the three consecutive stable baseline sessions were combined for each individual rat to allow sufficient trials numbers for model fitting. Nevertheless, not all rats completed sufficient trials of each outcome type to allow for satisfactory model fitting, and so were excluded. The numbers of animals used for this analysis were: males: n = 41; females: n = 63.

### Cluster analysis

To investigate whether workers and slackers were distinct groups, K-means clustering (using the kmeans function in Matlab) was carried out using data from all six cohorts averaged across the three stable baseline sessions after acquisition. Behavioral measures (% HR choice, % choice omissions, % premature responses, % HR accuracy and % LR accuracy) and model parameters (drift rate, starting point, boundary and non-decision time) were included. The average of three baseline sessions from the week immediately preceding the beginning of a drug study were used to get accurate classifications for reanalysis of each pharmacological manipulation. The optimal number of clusters was determined using the evalclusters function in Matlab, with the Silouhette value as a criterion, and confirmed using evalclusters with the Gap Value criterion. These clusters were applied to analyse behavioral measures and model parameters from stable baseline sessions, as well as drug studies.

### Statistics

Behavioral measures and model parameters were analysed with mixed ANOVAs with sex (female or male) and cluster as between-subjects factors, and session as the within-subjects factor. Differences in model parameters and behaviors between clusters were analyzed using one-way ANOVAS, with Tukey’s HSD for post-hoc comparisons as appropriate. For comparison with previous analyses, behavioral measures and model parameters were also analysed using group (worker or slacker) instead of cluster.

For reanalysis of drug studies, mixed ANOVAs with cluster as between-subjects factor and dose as the within-subjects factor were carried out for behavioral measures (% HR choice, HR choice latency, LR choice latency, % omissions, % premature responses, % HR accuracy and % LR accuracy) and model parameters. Bonferroni corrected pairwise comparisons were performed as post-hoc tests as appropriate. Huynh-Feldt corrections were used to adjust for violations of the sphericity assumption. Statistics are reported with the ANOVA F-value (degrees of freedom, error) and p-value as well as any post-hoc p-values. All statistical analysis was conducted using SPSS 28.0.0.0 for Windows (IBM SPSS Statistics) or Matlab R2021b for Windows (The Mathworks), and all graphs were produced using GraphPad Prism 9.4.0 for Windows (Graphpad Software, USA).

## Results

### Cluster analysis

Using behavioral measures (HR choice, choice omissions, premature responses, HR accuracy, and LR accuracy) combined with the four main diffusion model parameters (zr, a, v and t0) from stable baseline rCET data, it was determined that four clusters were optimal for this data (Silhoutte value = 0.3095, compared with 0.2406, 0.2308 and 0.2449 for 2, 3 and 5 clusters respectively). K-means analysis with four clusters grouped rats similarly to the previously used metric (70% choice of HR trials), and based on the results we interpreted these clusters as representing: “good workers”, “poor workers”, “good slackers” and “poor slackers”. Only 7 out of the 157 total individual rats in the analysis were assigned to the opposite group type (either worker or slacker) in the cluster analysis compared to the traditional method (Table 2). Table 2 also contains a breakdown of the numbers of males and females in each cluster.

**Table 2-.**
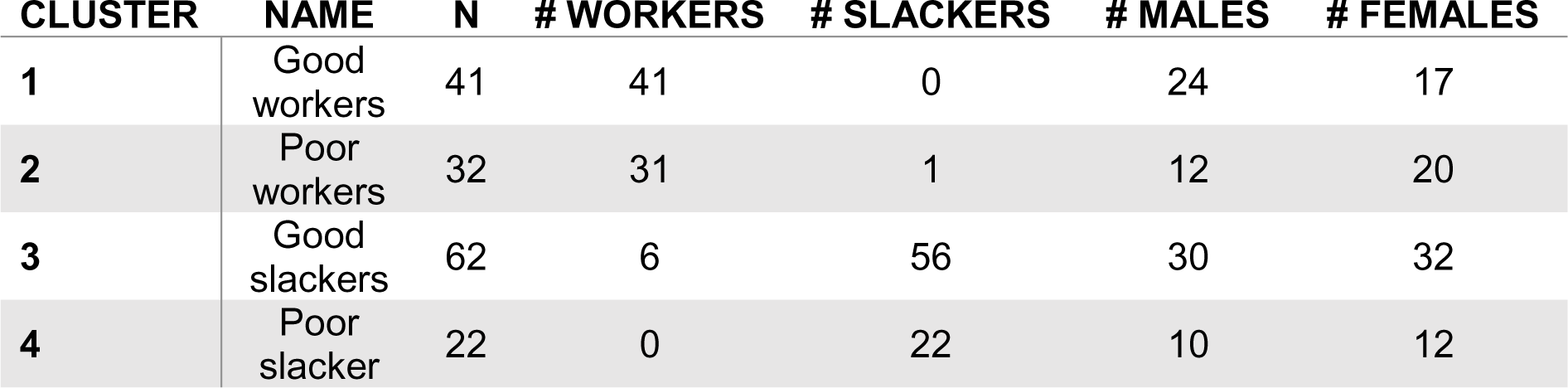
Numbers of rats for each of the four clusters from the k-means cluster analysis.

| **CLUSTER** | **NAME** | **N** | **# WORKERS** | **# SLACKERS** | **# MALES** | **# FEMALES** |
| --- | --- | --- | --- | --- | --- | --- |
| **1** | Good workers | 41 | 41 | 0 | 24 | 17 |
| **2** | Poor workers | 32 | 31 | 1 | 12 | 20 |
| **3** | Good slackers | 62 | 6 | 56 | 30 | 32 |
| **4** | Poor slacker | 22 | 0 | 22 | 10 | 12 |

### Analysis of stable baseline data: main analysis

#### Behavioral data analyzed by cluster

There were no cluster or group by sex interactions, so behavioral measures are presented collapsed by sex (Figure 2). The good worker cluster comprised rats that picked the HR choice most often (85.27±7.18%), and significantly more so than the two slacker clusters (cluster: F_3,149_=238.115, p<0.001, Tukey’s HSD: p’s<0.001). This cluster had the best performance across other behavioral measures, in that they made the fewest choice omissions (3.43±3.07%) and least premature responses (6.06±3.85%), as well as being significantly less premature than poor workers (cluster: F_3,149_=5.148. p=0.002, Tukey’s HSD: p=0.006). Good workers were most accurate after HR choices (75.10±5.93%), significantly more so than all other clusters (cluster: F_3,149_=24.396, p<0.001; Tukey’s HSD: p’s<0.001), and most accurate for LR choices (94.72±3.93%), although only significantly so compared to good slackers (cluster: F_3,149_=33.616, p<0.001, Tukey’s HSD: p=0.035). Poor workers made numerically (82.71±7.83%), but not significantly fewer HR choices than good workers, but performed the worst on other behavioral measures. These rats made numerically most omissions (5.43±6.84%) and premature responses (9.51±4.82%), were significantly more premature than good workers (p=0.006), had the lowest HR accuracy (59.87±5.89%), significantly lower than good workers (p<0.001) and good slackers (p<0.001), and were the worst at detecting the stimulus after LR choices (83.64±6.85%; p’s<0.001 for post-hoc comparisons to all other clusters). There were no differences in choice latencies between clusters (F’s≤2.490, p’s≥0.063).

**Fig. 2-.**
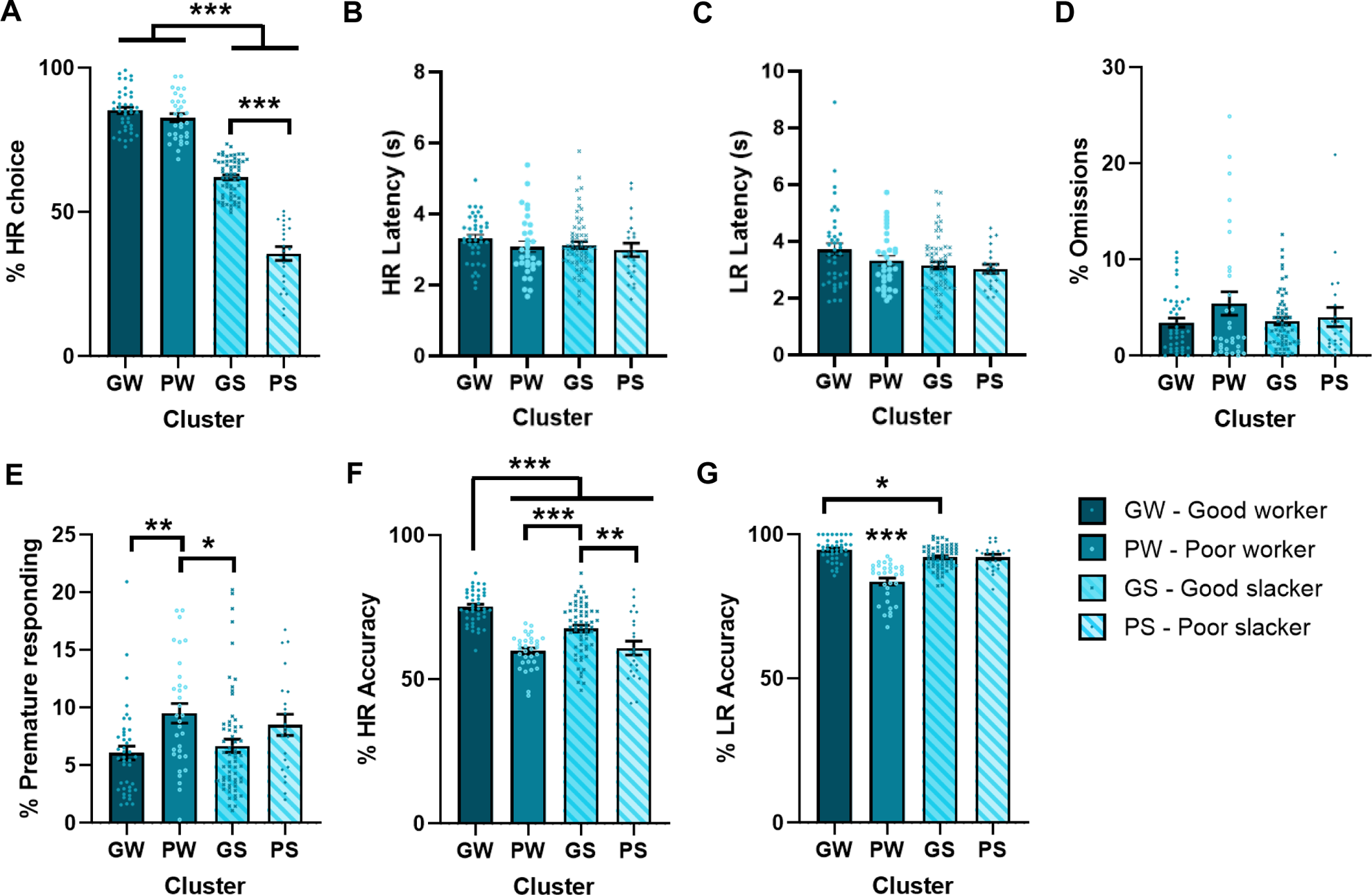
Behavioral measures for the four clusters identified by k-means clustering. K-means clustering with four clusters revealed four distinct groups, which we labelled as good workers (GW), poor workers (PW), good slackers (GS) and poor slackers (PS). (A) The two worker clusters make more HR choices than the two slacker clusters, with poor slackers choosing HR trials the least. (B/C/D) There were no significant differences between clusters in response latencies for HR or LR choices, or percentage of choice omissions. (E) Poor workers make more premature responses than good workers or good slackers. (F) Good workers were most accurate to detect the stimulus on HR trials, followed by good slackers. Poor workers were the least accurate. (G) Poor workers were less accurate than all other clusters on LR trials. Good workers were more accurate than good slackers. Good workers: n = 41; poor workers: n = 33; good slackers: n = 62; poor slackers: n = 22;. Bars are mean ± SEM with individual data points laid over the top. *p<0.05, **p<0.01, ***p<0.001.

Good slackers made more HR choices than poor slackers (62.04±6.56%; Tukey’s HSD: p<0.001), but fewer HR choices than either worker cluster (p’s<0.001). There were no significant differences between good and poor slackers in choice omissions, premature responses, or LR accuracy, although numerically good slackers made fewer omissions (3.58±2.96% compared to 4.02±4.66% for poor slackers) and fewer premature responses (6.68±4.50% compared to 8.52±4.32%). Good slackers were more accurate on HR trials compared to poor slackers (67.60±9.08% versus 60.85±11.35%, Tukey’s HSD: p=0.004), less accurate than good workers (p<0.001) but more accurate than poor workers (p<0.001).

#### Behavioral data analyzed by group

We also analysed this large dataset using traditional worker/slacker groupings for comparison to published work (Figure S1). Contrary to previous reports (Cocker et al., 2012; Hosking et al., 2014; 2015; 2016; Silveira et al., 2017; 2018; 2020; 2021), workers were more accurate on HR trials, and less accurate on LR trials (HR accuracy: group: F_1,153_=11.549, p<0.001; LR accuracy: F_1,152_=4.395, p=0.038). The small size of these differences (2-3%) likely explains why these failed to reach significance when each smaller cohort was analysed separately (HR: workers: 68.69±0.70%, slackers: 65.41±0.83%; LR: workers: 89.93±0.75%; slackers: 91.81±0.41%). Workers were also slower to choose their less-preferred LR option (group: F_1,149_=17.673, p<0.001). There were no group differences in any other behavioral variables analysed, as reported previously (group: all F’s≤1.949, all p’s≥0.144).

#### Comparison of task performance between males and females

Males were more accurate on HR trials (sex: F_1,149_=9.909, p=0.002) and made fewer choice omissions (sex: F_1,149_=6.100, p=0.015, Figure 3). No other variables were significantly different across the sexes (group: all F’s≤2.043, all p’s≥0.155). Numbers of males and females in each cluster are shown in Table 2. There were approximately equal numbers in the two slacker clusters, with slightly more male good workers, and slightly more female poor workers. Using traditional criterion, workers and slackers were almost equally split across males and females (Table 1, and behavioral data grouped by worker/slacker groupings are shown in Figure S2).

**Fig. 3-.**
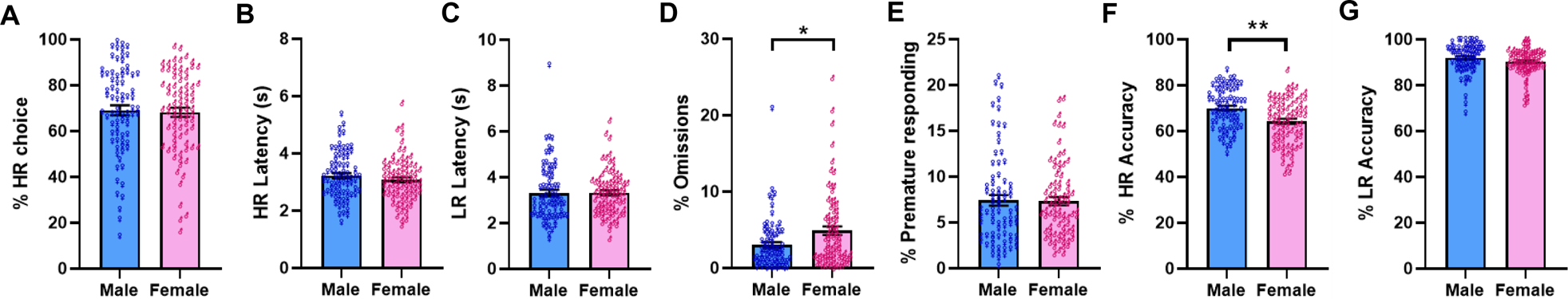
Comparison of task performance between males and females. (A-C) There were no differences between males and females for percentage HR choices, or latencies to make a HR or LR choice. (D) Males make fewer choice omissions than females. (E) There were no sex differences in premature responding. (F) Females have lower accuracy to detect the stimulus after a HR choice, but (G) there were no differences for LR accuracy. Males: n = 76; females: n = 81. Bars are mean ± SEM with individual data points laid over the top. *p<0.05, **p<0.01.

#### Model parameters analyzed by cluster

There were no cluster or group by sex interactions for model parameters, so data is presented collapsed by sex (Figure 4). Good workers had neutral decision starting points (0.53±0.14, one sample t-test compared to a value of 0.5: p=0.175), the widest decision boundaries (4.60±0.80), but only significantly wider than the two slacker clusters (F_3,149_=19.157, p<0.001, Tukey’s HSD: p’s<0.001), steeper drift rates towards HR choices than all other clusters (0.38±0.18; F_3,149_=26.020, p<0.001; Tukey’s HSD: p’s<0.03), and the shortest non-decision time (0.33±0.18), significantly shorter than poor slackers (F_3,149_=2.825, p=0.041, Tukey’s HSD: p=0.021). Poor workers had positive decision starting points (0.66±0.15, one sample-test versus 0.5: p<0.001) that were different from all other clusters (p’s≤0.013) and wide decision boundaries (4.44±0.90) that were wider than both slacker clusters (p’s<0.001). They had moderate drift rates (0.25±0.17) that were more negative than good workers (p=0.030) and more positive than poor slackers (p<0.001), and moderate non-decision times (0.40±0.26).

**Fig. 4-.**
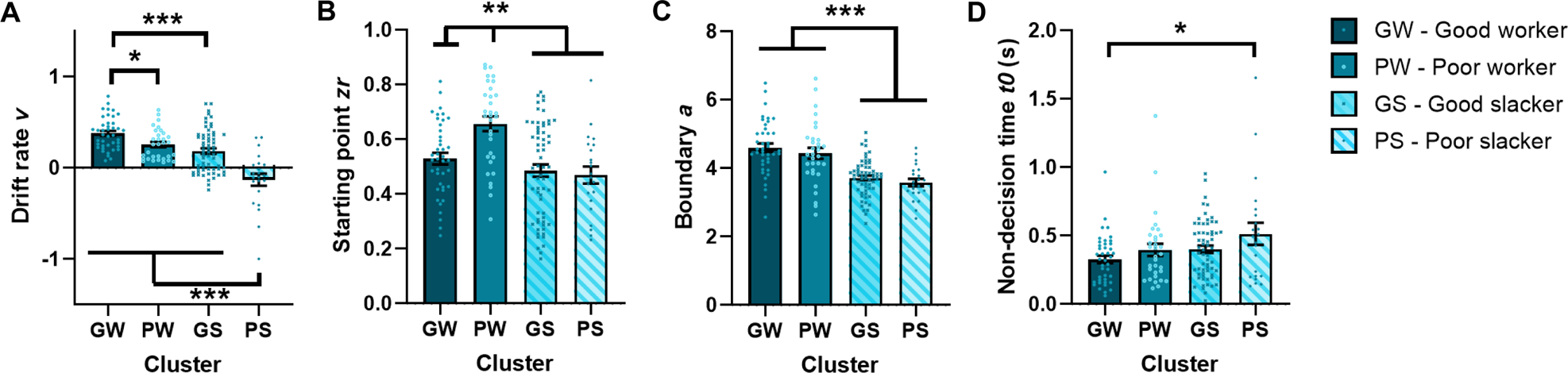
Model parameters for the four clusters identified by k-means clustering. K-means clustering with four clusters revealed four distinct groups, which we labelled as good workers (GW), poor workers (PW), good slackers (GS) and poor slackers (PS). (A) Good workers had the most positive drift rates and poor slackers had the least positive drift rates. Poor workers and good slackers did not differ. (B) Poor workers had positively biased decision starting points that were different from all other clusters, which had neutral starting points. (C) Boundaries were wider for the two worker clusters compared to the two slacker clusters. (D) Non-decision times were longer in poor slackers. Good workers: n = 41; poor workers: n = 33; good slackers: n = 62; poor slackers: n = 22;. Bars are mean ± SEM with individual data points laid over the top. *p<0.05, **p<0.01, ***p<0.001.

In both slacker clusters these behavioral profiles corresponded to neutral decision making starting points (0.48±0.18 and 0.47±0.15 for good and poor slackers respectively) and smaller distances between decision boundaries as compared to both worker clusters (good slackers: 3.71±0.53, poor slackers: 3.58±0.51, Tukey’s HSD compared to worker clusters: p’s<0.001). Good slackers had more positive drift rates than poor slackers (0.18±0.24 versus −0.13±0.31; p<0.001), and poor slackers had the longest non-decision times (0.51±0.38), significantly so compared to good workers only (p=0.021).

#### Model parameters analyzed by group

Using traditional groupings (Figure S3), workers had more positive drift rates than slackers (group: F_1,153_=39.468, p<0.001), but also had decision starting points that were biased towards the HR boundary, unlike slackers that had neutral starting points (group: F_1,153_=13.071, p<0.001, one sample t-tests against a theoretical value of 0.5 for workers: p<0.001). Boundary separation was greater for workers than slackers (group: F_1,153_=46.790, p<0.001). Non-decision time did not differ between groups.

#### Comparison of model parameters between males and females

As shown in Figure S3, males had more positive drift rates than females (sex: F_1,153_=5.840, p=0.017) and neutral starting points whereas starting points for females were biased towards the HR boundary (sex: F_1,153_=5.157, p=0.025, females: one sample t-tests against a theoretical value of 0.5: p=0.008). Boundary separation and non-decision time did not differ between the sexes. Data split by sex and group are shown in Figure S4.

### Analysis of stable baseline data: outcome analysis

#### Relationship between DDM parameters and trial type (correct, incorrect, premature) analyzed by cluster

There were no interactions between outcome or cluster and sex, and so data is presented for males and females combined (Figure 5). There was a main effect of sex for starting point (F_1,96_=4.918, p=0.029), recapitulating the finding from the main analyses whereby females have more positive starting points than males (Figure S5). There were no other sex specific effects.

**Fig. 5-.**
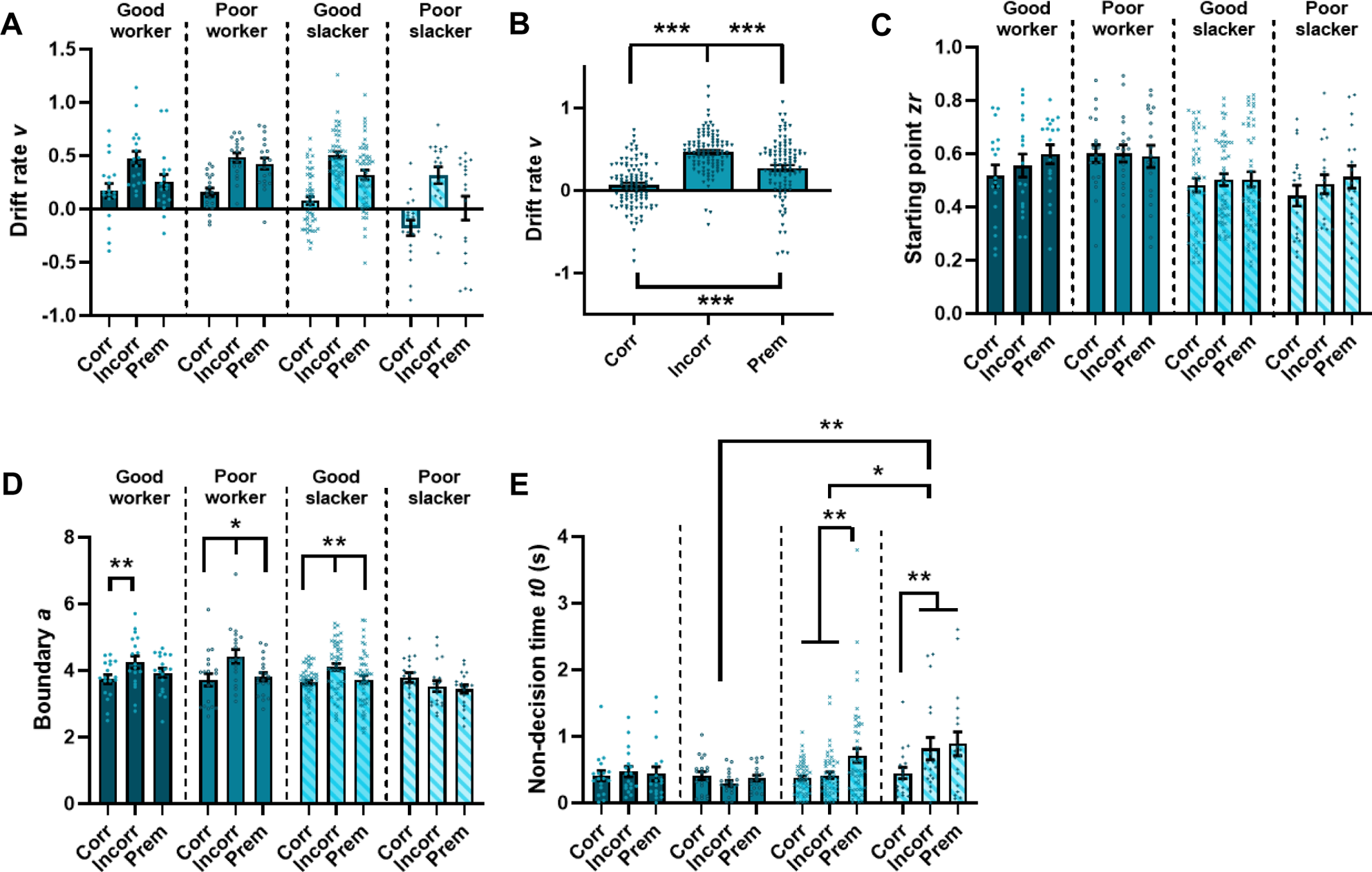
Diffusion modeling of baseline behavioral data from the rCET – outcome condition. Diffusion model parameters (zr, a, v, t0) fit to data from three stable baseline sessions combined from the rCET, split into trials where the rat went on to make a correct, incorrect or premature response after the choice. (A) Drift rate differs for each outcome, but not by cluster, so data was collapsed across clusters as shown in (B). (B) Drift rates were steepest for incorrect responses, smallest for correct responses and intermediate for premature responses. (C) Starting point did not vary by outcome. (D) Boundaries are wider for incorrect responses in both worker clusters, and for good slackers, but not different in poor slackers. (E) Non-decision times are longer on trials followed by a premature response in both slacker clusters, as well as after an incorrect response in poor slackers. Workers: n = 37; slackers: n = 67; males: n = 41; females: n = 63. Bars are mean ± SEM with individual data points laid over the top. *p<0.05, **p<0.01, ***p<0.001.

Drift rate, boundary separation, and non-decision time all varied significantly by trial type (outcome: - drift rate: F_2.000,192.00_=62.751, p<0.001, - boundary separation: F_1.969,188.982_=11.658, p<0.001, - non-decision time: F_1.841,176.750_=7.013, p=0.002) and these patterns were significantly different across cluster for boundary and non-decision time (outcome*group: - boundary separation: F_5.906,188.982_=3.537, p = 0.003, - non-decision time: F_5.523,176.750_=2.521, p=0.027). Across all clusters, choices followed by a correct response had the least positive drift rates compared to other types, those followed by a premature response had intermediate drift rates, and incorrect trials the most positive drift rates (all comparisons: p’s<0.001; Figure 5B). Boundaries were not different between outcomes for poor slackers. For all other clusters, choices followed by an incorrect trial had wider boundaries than those followed by a correct trial (p’s≤0.002), and were also wider than premature trials in the poor worker (p=0.031) and good slacker (p=0.004) clusters. Non-decision times were longer for choices followed by a premature response compared to a correct one (p=0.003), and this was driven by differences in both slacker clusters (p’s≤0.006). The poor slacker cluster had non-decision times for incorrect outcome trials that were also longer than the good slacker (p=0.014) and poor worker (p=0.003) clusters.

#### Relationship between DDM parameters and trial type (correct, incorrect, premature) analyzed by worker/slacker grouping

Similar to the cluster analysis, all parameters except starting point were different between outcomes (outcome: - drift rate: F_1.915, 191.500_=65.377, p<0.001, - boundary separation: F_2,200_ =19.513, p<0.001, - non-decision time: F_1.831,183.103_=7.367, p=0.001; Figure S5). Drift rate and boundary varied by worker or slacker group (outcome*group: - drift rate: F_1.915,191.500_=5.873, p=0.004; - boundary: F_2,200_=3.355, p=0.039). Choices followed by a correct response had the least positive drift rate compared to other trial types, a pattern that held across workers and slackers (p’s<0.001), but the drift rate was relatively more positive in workers than slackers on correct trials (p<0.001).

Choices followed by a premature response had intermediate drift rates that were significantly different from those followed by a correct or incorrect response (p’s≤0.002). Workers again had more positive drift rates than slackers on these trials (p=0.005). Non-decision times were significantly longer on these premature trials (ps≤0.014).

Choices followed by an incorrect response had the most positive drift rate, an effect that was consistent across workers and slackers (p’s<0.001). These choices also had the widest boundary separation (p’s≤0.003). This distinction was most pronounced in workers (p=0.003), and boundaries were generally wider across all trial types in these animals (group: F_1,102_=4.153, p=0.044), replicating the finding from the main analysis.

### Re-analysis of drug challenge data

#### 1) Serotonergic drug manipulations (Silveira et al. 2020)

**Ro-60-0175:** In previous analyses, Ro-60-0175 had no effect on preference for HR trials, or on choice latencies, but the 0.1 and 0.6 mg/kg doses increased HR accuracy. Reanalyzing by cluster (Figure S6) confirmed that the increase in HR accuracy was common across all animals (no dose*cluster interaction: F_9,54_=0.237, p=0.987, post-hoc: 0.1 mg/kg: p=0.018; 0.6 mg/kg: p=0.004), but did reveal that 0.3 and 0.6 mg/kg doses of Ro-60-0175 specifically increased choice omissions in poor slackers only (dose*cluster: F_9,54_=2.259, p=0.032, pairwise comparisons: p’s≤0.033). Ro-60-1075 did not cause any changes to model parameters (Table 3), as although there was a dose*cluster interaction for non-decision time (F_9,54_=2.130, p=0.042), there were no significant changes after correction for multiple comparisons.

**Table 3-.**
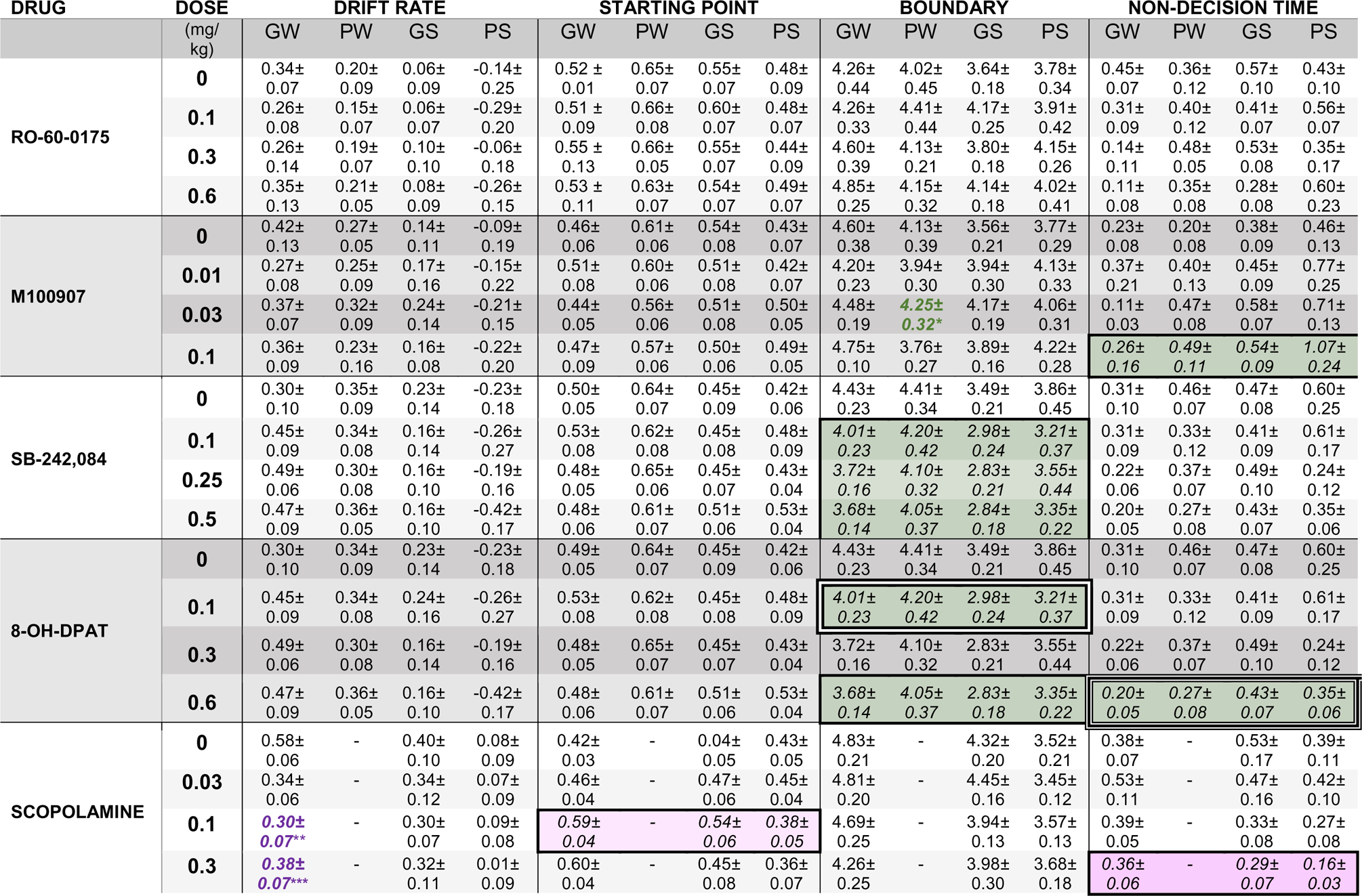
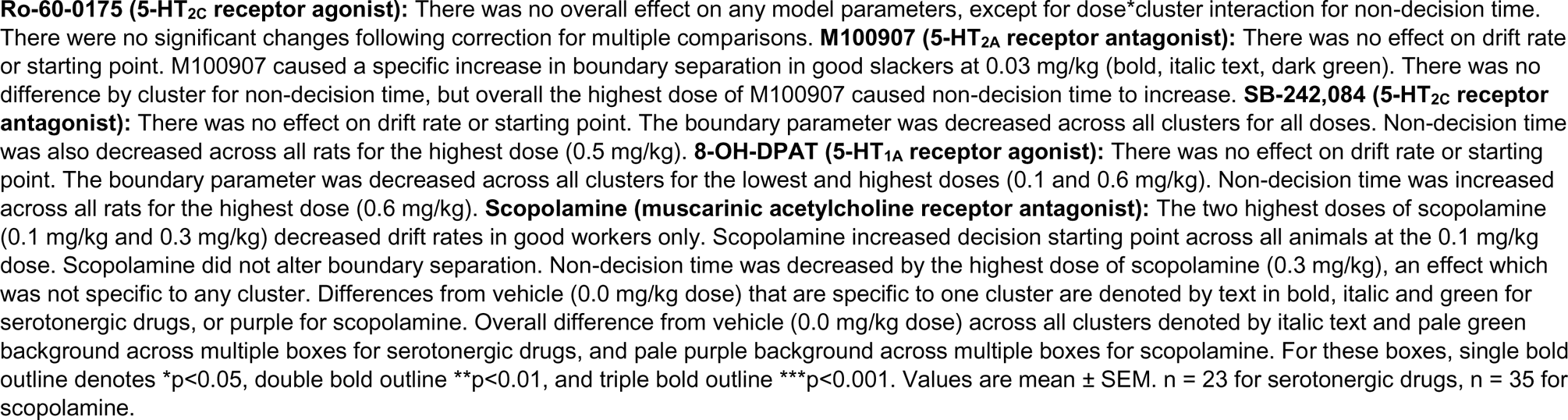
Diffusion model parameters fit to rCET data following administration of serotonergic drugs and scopolamine.

**M100907:** Previously, the two highest M100907 doses (0.03 and 0.1 mg/kg) increased RTs for both HR and LR choices, and also increased choice omissions. All doses decreased premature responses on HR trials. Reanalysis using clusters (Figure S7) found that the changes for latencies and choice omissions were common to all clusters (no interactions: F’s≤1.500, p’s≥0.170), but the drug’s effect on premature responses varied by cluster (dose*cluster: F_7.986,50.579_=2.132, p=0.050). The decrease in premature responses was driven by both slacker clusters, with significant reductions at 0.01 mg/kg in good slackers (p=0.039), and 0.1 mg/kg in poor slackers (p=0.049). This reanalysis also showed that the highest dose of M100907 caused a decreased in LR accuracy in poor workers only (dose*cluster: F_9,57_=2.789, p=0.009; Tukey’s HSD: p=0.022). Modeling revealed (Table 3) that the increased RTs were explained by an increase in the non-decision time across all clusters (dose: F_3,57_=4.498, p=0.007; pairwise comparisons: 0.0 vs 0.1: p=0.008), as well as a specific increase in boundary in good slackers for the 0.03 mg/kg dose (dose*cluster: F_3,57_=2368, p=0.013, Tukey’s HSD: p=0.013).

**SB 242,084:** In the prior analyses, SB 242,084 decreased RTs for both HR and LR trials, and also increased premature responses across both trial types. Again, changes in latencies were common across clusters in the reanalysis (Figure S8; no interactions: F’s≤1.517, p’s≥0.16), but varied by cluster for premature responding (dose*cluster: F_9,57_=2.672, p=0.012). SB 242,084 did not alter premature responses in good workers, but did increase this measure in all other clusters (p’s≤0.035). In this case, the change in RTs for HR and LR trials was explained by both reduced boundaries (Table 3; dose: F_3,57_=15.594, p<0.001, Tukey’s HSD for all doses: p’s<0.001) and decreased non-decision times (dose: F_3,57_=3.122, p=0.033; pairwise comparisons: 0.0 vs 0.25: p=0.047, 0.0 vs 0.5: p=0.016).

**8-OH-DPAT:** In the original analyses, the highest dose of 8-OH-DPAT (0.6 mg/kg) increased the latency to make a LR choice, increased choice omissions, and also impaired HR accuracy. None of these changes were specific to any clusters (Figure S9; no interactions: F’s≤1.389, p’s≥0.215). There were two model parameter changes, again not specific to cluster (Table 3). The boundary was decreased for both the 0.1 and 0.6 mg/kg doses (dose: F_3,57_=4.110, p=0.010; Tukey’s HSD: 0.0 vs 0.1 mg/kg: p=0.008; 0.0 vs 0.6 mg/kg: p=0.047), whilst non-decision time was increased for the highest dose only (dose: F_3,57_=13.842, p<0.001; pairwise comparisons: 0.0 vs 0.6 mg/kg: p<0.001). This suggests that non-decision time was driving the increase in latency to make a LR choice, as the decreased boundary was not specific to the highest dose, and the change in boundary was in the opposite direction to the change in RT.

#### 2) Scopolamine (Silveira, 2018)

In previous analyses, scopolamine decreased preference for the HR option across all rats at the highest dose only, but when analyzed by group this difference was driven by workers. Scopolamine also decreased HR choice latencies across all doses while also increasing choice omissions. For this reanalysis (Figure S10), no animals were classified as poor workers. Reanalysis revealed that choice latencies and omissions did not vary by cluster (no interactions: F’s≤1.664, p’s≥0.138), but the reduction in HR choice was driven purely by good workers (dose*cluster: F_2.449,78.359_, p=0.001, pairwise comparison: p=0.009). This was explained by less steep drift rates in good workers (Table 3; dose*cluster: F_5.365,85.846=2.613_, p=0.027, pairwise comparisons: 0.0 vs 0.1: p<0.001; 0.0 vs 0.3: p=0.007). The decrease in HR choice latencies was explained by a combination of the starting point moving closer to the HR boundary (dose: F_2.684,85.891_=3.797, p=0.016; post-hoc: 0.0 vs 0.1: p=0.035), along with a decrease in the non-decision time (dose: F_2.800,89.595_=3.310, p=0.026; post-hoc: 0.0 vs 0.3: p=0.015), effects that occurred across all clusters. The boundary parameter was not altered by scopolamine, but was greater in good workers compared to poor slackers across all doses, similar to results from the larger cohort (cluster: F_1,32_=6.502, p=0.004; Tukey’s HSD: p=0.001).

## Discussion

In this study we used the drift diffusion model combined with cluster analysis to analyze rodent behavioral data from the rCET to provide insight into the decision making processes underlying choices based on evaluation of cognitive effort. This analysis framework revealed that, broadly speaking, the use of traditional worker/slacker groupings based on a median split does offer a useful way to analyze rCET data, but that there is more nuance within these populations. Overall, the differences between performance of workers, those rats that choose the more effortful but more highly rewarded option most frequently, and slackers (the converse) is explained by workers having steeper rates of evidence accumulation, increased distance between thresholds and shorter non-decision times. Clustering analysis revealed that within these worker and slacker classifications, rats are using different strategies to perform the task. A subset of workers have increased rates of information accumulation with no a-priori bias towards hard choices and perform better overall on the task, whilst the other subset have less strong rates of information accumulation and their decisions begin closer to the hard choice boundary. These workers actually perform the worst on the attentional part of the task of all rats. The two slacker groups are more similar to each other. “Good slackers” choose the hard choice more often (but less so than workers), a preference reflected in the drift rate, and also show improved accuracy following hard choice trials. The other slacker group pick the hard choice very infrequently, have long non-decision times, and have low accuracy on the few hard choice trials they do choose, but show equivalent performance to good slackers on all other measures.

There were differences in decision making between males and females: males have no priori bias towards either decision but steeper rates of information accumulation, compared to females that have decision starting points biased towards hard choices along with less steep evidence accumulation. Modeling also revealed differences in the way that drug manipulations alter these processes. The behavioral dissociation seen between the effects of serotonergic drugs, which do not alter choice behavior but do effect accuracy and premature responding, and scopolamine, which has no effect on the former but does alter choice of the hard option, is further supported by modeling results. Broadly, changes in the boundary separation and non-decision time underlie serotonergic drug effects on HR/LR choice latencies across all rats, whilst differences in the drift rate and decision starting point explain scopolamine’s effects on both latency and number of HR choices made, with HR choice specifically driven by effects in good workers only.

Previous studies have demonstrated that some manipulations have opposing effects on worker and slacker performance, whilst others only change behavior in one or other group. For example, amphetamine reduced the choice of HR trials in workers, but increased them in slackers (Cocker et al., 2012), the same effect as seen with inactivation of the basolateral amygdala (Hosking et al., 2014). Caffeine reduced HR trial choice in workers only (Cocker et al., 2012), whilst nicotine reduced HR trial choice in slackers only (Hosking et al., 2012). Administration of cannabidiol partially rescued the reduction in HR trials caused by Δ9-tetrahydrocannabinol, but again only in slackers (Silveira et al., 2017). In all of these cases, differences between workers and slackers could not be explained by other factors, such as manipulations causing changes in motivation, attentional impairments, or inability to remember or associate the lever with its correct choice and reward value pairings (Cocker et al., 2012; Hosking et al., 2014; Silveira et al., 2017).

In the first rCET study, the authors proposed that differences between workers and slackers may be due to differing sensitivities to cognitive effort costs, or that animals use different strategies to perform the task, allowing some to more easily overcome cognitive demand (Cocker et al., 2012). In terms of the diffusion model, increased sensitivity to effort cost could manifest as a reduction in the starting point, i.e. slackers would be biased away from the HR trial boundary, or as a reduced drift rate, i.e. less clear integration of information leading to a choice for a HR trial. Diffusion modeling revealed that all four clusters differ in the gradient of the drift rate. Given that workers and slackers are classified using the same behavioral measure (number of choices for HR and LR trials) as is input into the model, it would be surprising if these groups did not have different drift rates. From plotting individual data points for the drift rate (Figure 4), considerable overlap between the four populations is evident, suggesting this is not the only factor explaining behavioral performance.

Both worker clusters chose the hard choice frequently, but a proportion of those rats carry on to perform well on the attentional part of the behavioral task (as well as making the fewest choice omissions). Conversely, the other proportion of workers perform the worst as indicated by the other task behavioral measures, with lowest accuracies to detect the stimulus, highest premature rates and the most choice omissions. These two subsets show distinct decision making profiles on the choice part of the task: good workers strongly integrate information to reach the hard choice boundary with no a priori bias for either option, whilst poor performing workers have decision starting points biased towards the hard choice boundary along with lower rates of evidence accumulation. In this subset of worker rats, performance is clearly linked with cognitive effort allocation, and suggests that they are more sensitive to the effort costs associated with the hard choice. Modeling indicates these rats employ a strategy which allows them to “more easily” choose the hard choice by having the decision start closer to the hard choice boundary so less effort is required to integrate information, but they then fail to perform well on the rest of the task. These rats may be biased towards selecting the HR option, perhaps because of the larger reward, but do not integrate their own cognitive performance or chances of success into the decision process.

It is interesting that the decision making profiles for males and females recapitulate the pattern seen in the cluster analysis for good and poor workers. Females rats on average have decision starting points biased towards a hard choice, with less steep rates of evidence accumulation, similar to poor workers, whilst modeling results for males more closely match good workers: neutral starting point but steeper information accumulation. Although there are relatively more males that are good workers and females that are poor workers (Table 2), these differences in numbers (7 and 8 in each cluster) are too small to account for all of the effect seen in the main analysis. It therefore seems that males and females employ different decision making strategies to perform cognitively effortful decisions, perhaps due to underlying differences in sensitivity to effort costs.

The two slacker clusters were much more similar to each other, with mostly overlapping values for behavioral measures and model parameters. Both slacker clusters have narrower boundaries than either worker cluster, suggesting that overall workers have a more cautious response profile, requiring more information to be accumulated before a decision is made. Between slacker clusters, slackers who very infrequently pick the hard choice (and perform with lower accuracy when they do) have longer non-decision times. This suggests that extraneous factors that are not directly related to integrating information regarding the decision have a bigger influence on their performance, potentially suggesting these rats might be more distractible and less focused, and therefore less driven to choose the harder trials. Together, these results provide clear evidence that rats are employing different strategies to perform the task, and that variability in the decision making process evaluating cognitively effortful options is reflected in attention-dependent accuracy and other performance metrics later in the task. As such, contrary to conclusions obtained from behavioral analyses alone, the processes involved in adjudication of cognitive effort are not completely independent from its deployment.

The diffusion model analysis split by trial outcome provides further insight into the differential decision making strategies used both within the task and between different rats. Drift rates showed the predicted pattern: drift rates were lowest (i.e. towards LR boundary) for trials followed by a correct choice, and this outcome occurs more often on LR trials. Similarly, drift rates were most positive (i.e. towards HR boundary) on incorrect trials, which are more frequent following HR choices. Drift rates for trials followed by a premature response (which occur with more similar frequency after both HR and LR choices) were in between these values. Boundaries were widest on trials followed by an incorrect response in all clusters apart from poor slackers, suggesting a possible trade-off between cognitive resources used during the choice vs visuospatial discrimination phases of the task. If more cognitive resources are used integrating information when deciding whether to select a HR or LR trial, this leaves less available for the attentional challenge immediately following the choice. As this effect was more pronounced in workers, it again suggests that depletion of cognitive resources during the evaluation of cognitive effort impacts other aspects of task performance. Non-decision time was longer in both good and poor slackers on choices preceding a premature response, as well as poor slackers for trials preceding an incorrect response. This further supports the interpretation that these rats may be more susceptible to extraneous, non-decision related factors, but that this specifically and negatively impacts their ability to inhibit responding (or respond correctly in poor slackers), after a choice.

For the serotonergic drug manipulations, modeling revealed that opposing behavioral effects were not necessarily explained by changes in the same model parameter in opposing directions. The serotonin system is complex, with more than 14 different receptor subtypes (Sharp & Barnes, 2020). Behaviorally, 5-HT_2A_ and 5-HT_2c_ receptors have been shown to have opposite functional effects, with 5-HT_2A_ receptor antagonism decreasing premature responding, whilst 5-HT_2c_ receptor antagonism increases this measure (e.g. Fletcher et al., 2007). On the rCET, the opposing effect on choice latency of the 5-HT_2A_ receptor antagonist M100907, and the 5-HT_2c_ receptor antagonist SB 242,084, were explained by changes in different model parameters: an increase in non-decision time for M100907 but a decrease in boundary for SB 242,084. The effects found in the model were observed at the same doses as had an effect on behavior, and across all rats.

8-OH-DPAT, a 5-HT_1A_ receptor agonist, also altered choice latency, but this increased for LR choices at the highest dose only. Modeling showed that both the boundary and non-decision time were altered at this dose. The decrease in boundary separation was not specific to the highest dose, so it is likely that increased non-decision time was driving the change in latency on LR trials. These results likely reflect subtle but important differences in the way that these serotonin receptor subtypes modulate neuronal function and therefore impact behavior, most likely due to their different pharmacology and receptor distribution across different brain regions (Sharp & Barnes, 2020).

Reanalysis using the cluster and model framework also exposed some behavioral changes in choice omissions, premature responding and accuracy that were specific to certain clusters. Ro-60-0175 specifically increased choice omissions in poor slackers only, despite no change in omissions being detected from the original analysis. The changes in premature responding caused by M100907 and SB 242,084 (which were found originally) were actually driven by both slacker clusters for M100907, and in all clusters except good workers for SB 242,084. M100907 also specifically decreased LR accuracy in poor workers only. These findings could suggest that certain decision making strategies are more easily disrupted by changes to serotonergic signalling. In future work, it would be interesting to investigate whether similar subpopulations with these types of decision profiles also exist in humans, and whether there is any link between these and differing susceptibilities for maladaptive decision making or risk for psychiatric disorders.

As noted previously, these associations between attentional task phase (5CSRT) performance and the decision-making process suggest that rats’ ability to successfully complete the visuospatial challenge is related to their decision-making strategies. However, it is important to note that we did not pinpoint any DDM parameter changes specifically associated with drug-induced increases or decreases in premature responding. While it is always possible that our modeling approach failed to represent every cognitive process happening during the choice phase of the task, it is also likely that behavioural inhibition can be regulated independently from the decision architecture, and through distinct neural circuits. Indeed, the 5-HT_2A/C_ ligands produce similar effects on premature responding on tasks which do not begin with a binary choice phase, such as the 5CSRT and rGT (Winstanley et al., 2003; 2004; Adams et al., 2017). Understanding how and when task complexity, and the resulting recruitment of additional cognitive processes, influences the behavioural effects of different pharmacological compounds remains largely understudied in rodents, and computational modeling could provide useful insight in this regard.

For the serotonergic drugs, which did not alter HR/LR choice, there were no changes in drift rate or starting point. In contrast, scopolamine, which decreased HR choices only in good workers and HR choice latencies across all animals, altered both of these parameters. The reduction in drift rate was specific to good workers, suggesting this explains the change in HR choice, whilst the increases in starting point (meaning there was a smaller distance to reach the upper boundary corresponding to a decision for the HR choice) and decreases in non-decision time were not specific to a particular cluster, and together may explain the reduced HR choice latency in all rats. Scopolamine’s effects on HR choice and drift rate being specific to good workers matches the initial rCET scopolamine study (Hosking et al., 2014). This result lends further support to the hypothesis put forward previously to explain differential effects on worker vs slacker performance, whereby there is an inverted-U relationship between cholinergic function and choice of HR trials, and workers and slackers sit to the left and right of the apex of the curve respectively. The modeling builds upon this by furthermore suggesting that the specific mechanism regulated by muscarinic cholinergic receptors is the evidence accumulation process. As such, antagonism of these receptors has a more negative impact on the rate of evidence accumulation in workers than in slackers, leading to reduced HR choice in the former group alone.

In sum, the use of the diffusion model and cluster analyses with these datasets has provided further insight into the decision making processes that underlie choices requiring cognitive effort in rats. Future studies could apply the same methodology to other drug or brain circuit manipulations that alter choice behavior on the rCET in order to further evaluate the utility of this approach. Although when analyzed at a single cohort level there are no obvious differences in behavioral measures capturing impulsivity or attention between workers and slackers, the use of modeling on a much larger dataset revealed different behavioral profiles that were linked to different decision making strategies, providing insight into why individual rats differ in their propensity to choose the HR reward. The current analysis suggests that choice behavior on the first stage of the task is not completely independent from attentional performance in the second phase, perhaps indicating that rats have a finite cognitive reserve to deploy on-task. Worker and slacker subgroups may differ either in the magnitude of this pool, or in the default allocations to different cognitive processes. A similar type of approach could be used with data from human studies to allow better translation and understanding of results from rodent studies with relevance to decision making perturbations that are seen across brain health and disease.

## Acknowledgements

This work was supported by a Discovery Grant awarded to CAW from the Natural Sciences and Engineering Research Council of Canada (NSERC; RGPIN-2017-05006). CAH was supported by a Michael Smith Health Research BC Trainee Award (#RT-2020-0564). MMS was supported by an NSERC Doctoral Award. BAH was supported by a Canadian Institutes for Health Research Doctoral Award. The experimental work took place at a UBC campus situated on the traditional, ancestral, and unceded land of the xʷməθkʷə^y̓^ əm (Musqueam), s^ə̓^ lílwətaʔɬSelilwitulh (Tsleil-Waututh) and Sḵwx̱ wú7mesh (Squamish) Peoples. We acknowledge and are grateful for their stewardship of this land for thousands of years.

