## Supplementary Figures for "Insight into differing decision-making strategies that underlie cognitively effort-based decision making using computational modeling in rats"

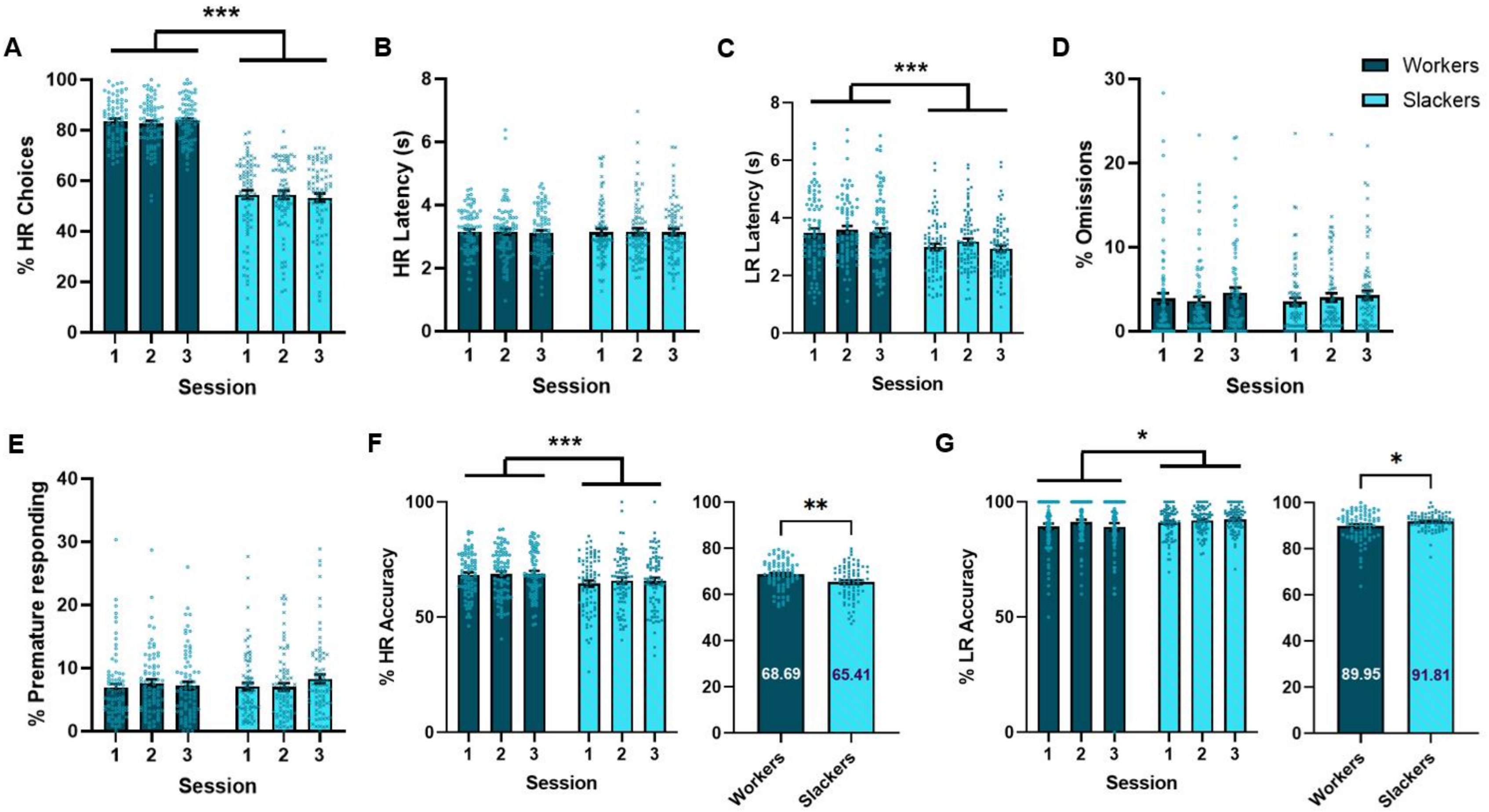


***Supplementary Figure 1 – Summarized behavioral data from three stable behavioral rCET sessions used for diffusion modeling analyzed by group***

Behavioral data from 6 cohorts of rats combined. (A) Workers made more HR choices than slackers. (B) Latency to make a HR choice was not different between groups, but (C) workers were slower to make an LR choice than slackers. (D-E) There were no differences in omission or premature responses. (F) Workers were more accurate to detect the stimulus on HR trials and (G) less accurate on LR trials. This is more clearly shown on the right graphs of both panels where data is collapsed across session. Workers: n = 78; slackers: n = 79; males: n = 76; females: n = 81. Bars are mean ± SEM with individual data points laid over the top. *p<0.05, ** p<0.01, *** p<0.001.

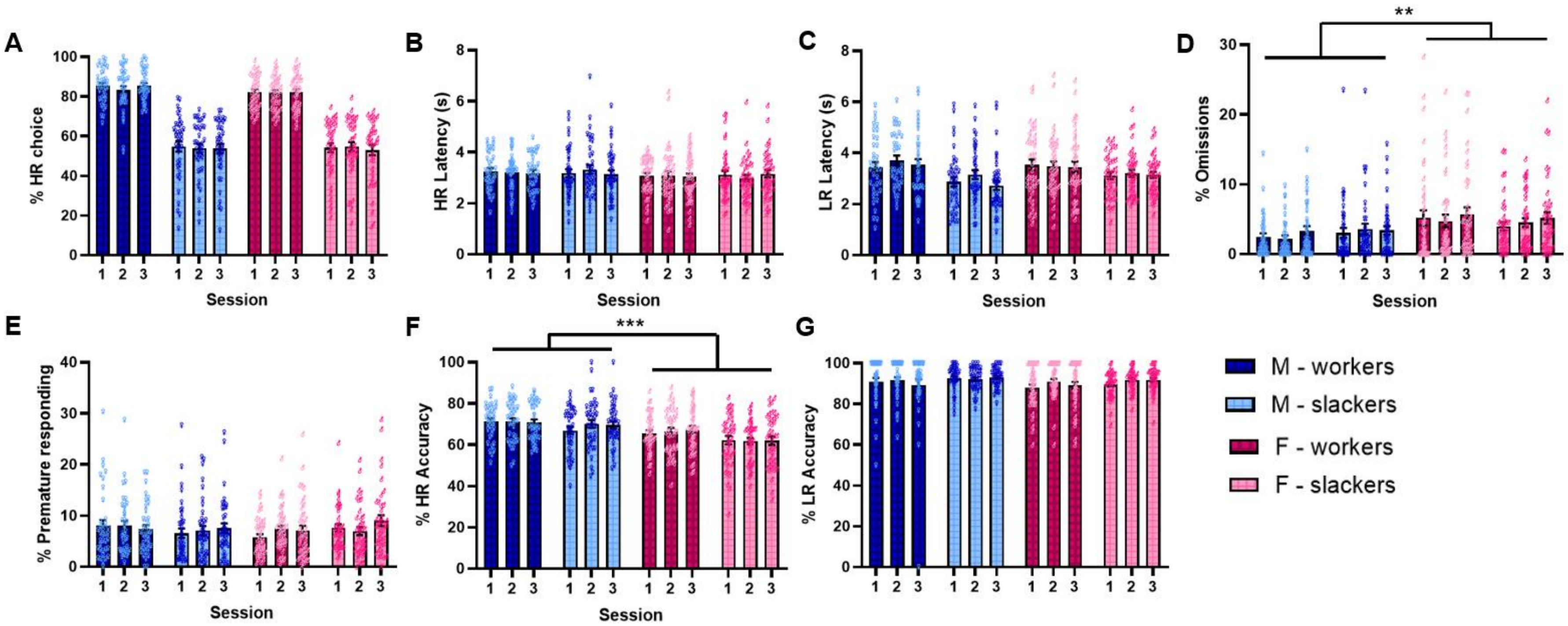


***Supplementary Figure 2 - Behavioral data from three stable behavioral rCET sessions split by sex.***

Behavioral data split by males and females for (A) percentage hard choices (HR) made in a session, choice latencies for (B) HR and (C) LR trials, (D) percentage of choice omissions, (E) percentage of premature responses after a choice, (F) accuracy to detect the stimulus following HR choices and (G) LR choices. (D) Females made more choice omissions than males, and (F) males were more accurate to detect the HR stimulus. Data from 6 cohorts of rats combined. Male workers: n = 37; male slackers: n = 39; female workers: n = 41; female slackers: n = 40. Bars are mean ± SEM with individual data points laid over the top.

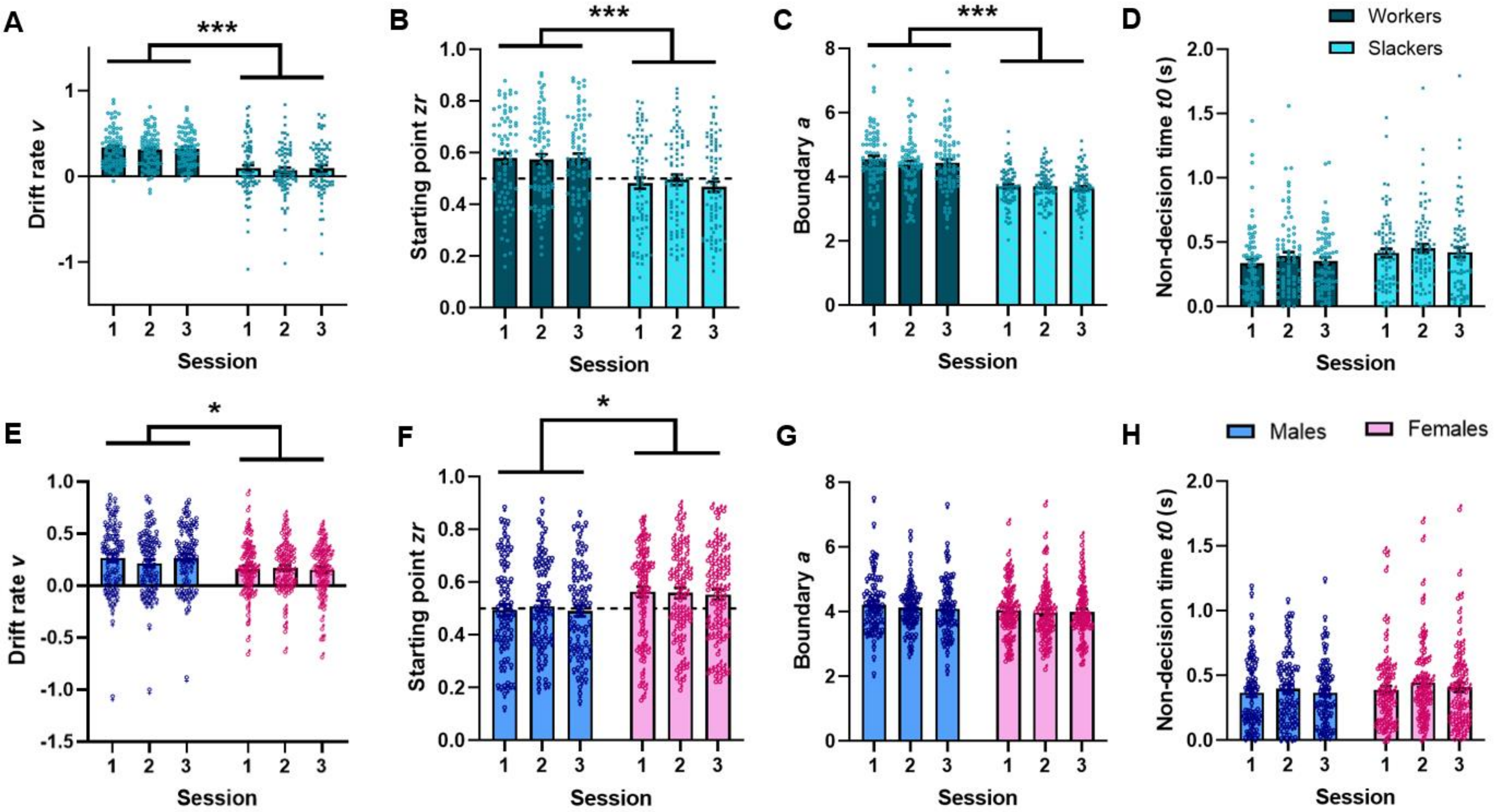


***Supplementary Figure 3 - Diffusion modeling of baseline behavioral data from the rCET – all trials condition analysed by group.***

Diffusion model parameters fit to three stable baseline sessions from the rCET, collapsed based on significant main effects (see Supplementary Figure 3 for full data). (A) Drift rate is greater for workers than slackers. (B) Workers have more positive starting points than slackers that are biased towards HR choices. (C) Workers have wider boundaries than slackers. (D) There was no difference in non-decision time. (E) Males have more positive drift rates than females. (F) Males have neutral starting points, while females have more positive starting points biased towards HR choices. (G-H) There were no sex differences for boundary or non-decision time. Workers: n = 78; slackers: n = 79; males: n = 76; females: n = 81. Bars are mean ± SEM with individual data points laid over the top. *$p<0.05$, ***$p<0.001$.

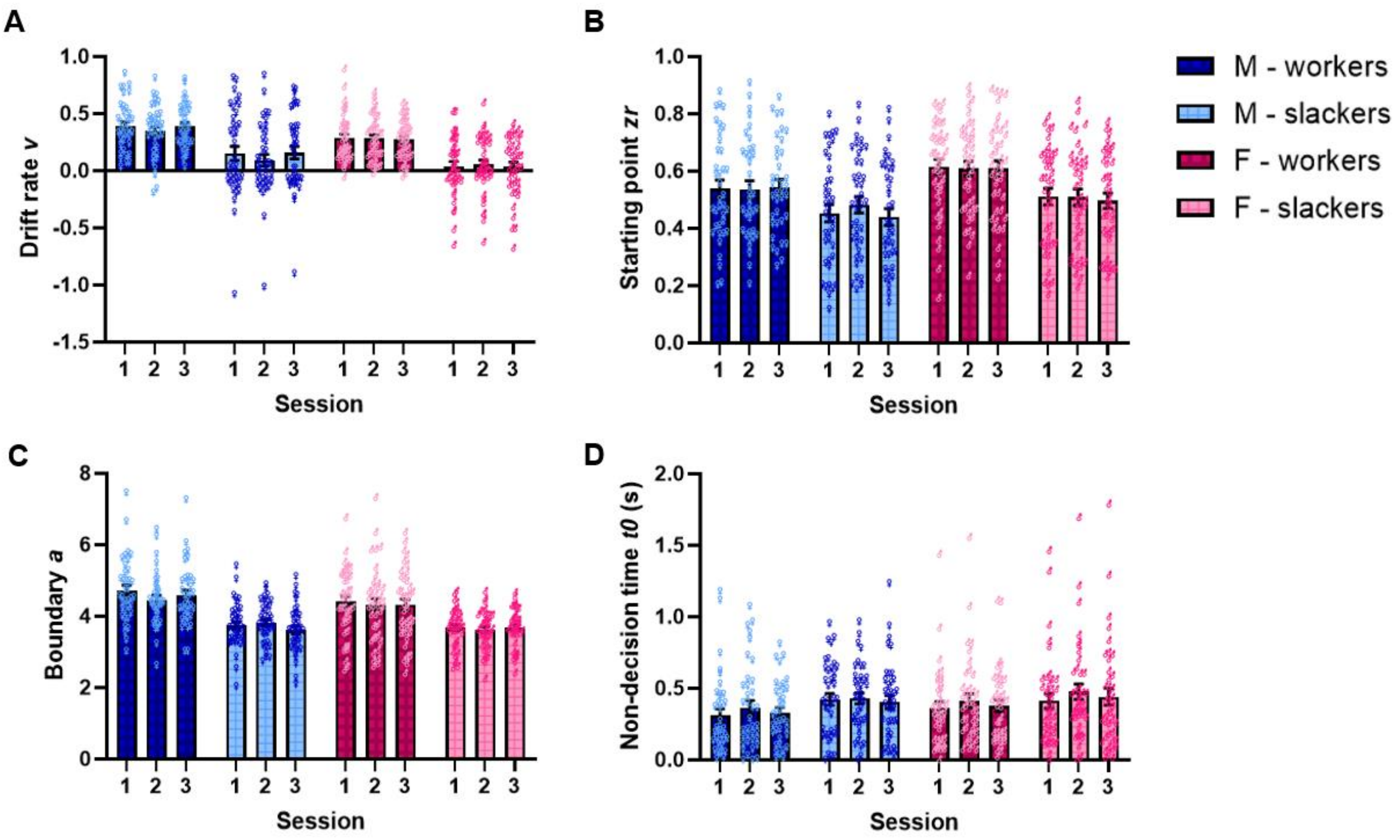


***Supplementary Figure 4 - Diffusion modeling of baseline behavioral data from the rCET – all trials condition split by sex and group.***

Diffusion model parameters fit to three stable baseline sessions from the rCET. Data from 6 cohorts of rats combined. Male workers: n = 37; male slackers: n = 39; female workers: n = 41; female slackers: n = 40. Bars are mean ± SEM with individual data points laid over the top.

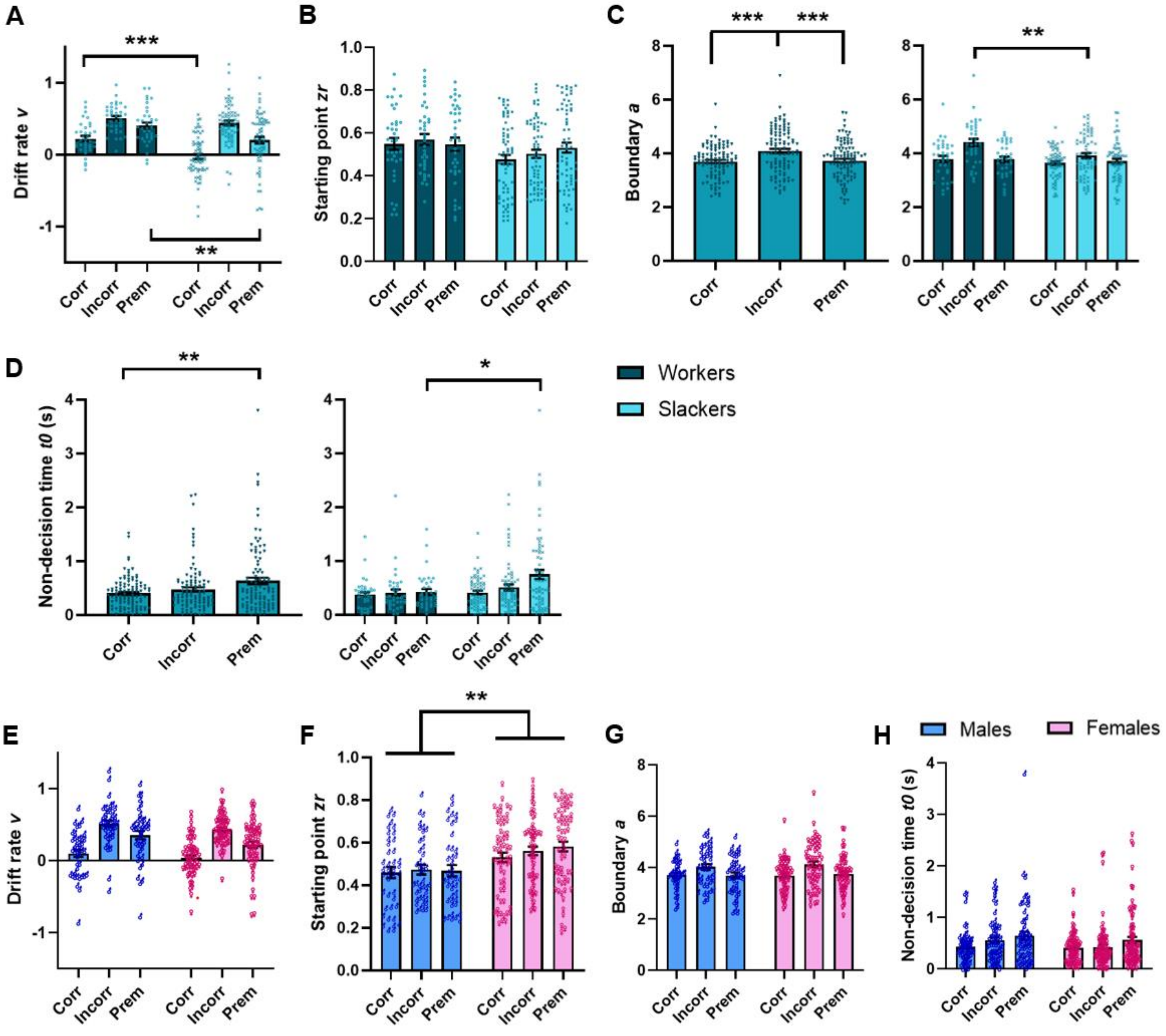

A
B
C
D
E
F
G
H
Drift rate v
Starting point zr
Boundary a
Non-decision time t0 (s)
Corr
Incorr
Prem
Workers
Slackers
Males
Females
***
**
*

***Supplementary Figure 5 - Diffusion modeling of baseline behavioral data from the rCET – outcome condition analyzed by group.***

Diffusion model parameters (zr, a, v, t0) fit to data from three stable baseline sessions combined from the rCET, split into trials where the rat went on to make a correct, incorrect or premature response after the choice. Data shown are collapsed based on significant main effects. (A) Drift rates are less steep in slackers compared to workers for choices made before a correct and premature response, but the same steepness between groups for choices made before an incorrect response. (B) Starting points did not differ by outcome. (C) Boundary is wider for incorrect responses, and this effect is amplified in workers compared to slackers. (D) Non-decision times are longer on trials followed by a premature response, driven purely by slackers. (E) Drift rates did not differ across outcomes between males and females. (F) Starting points are greater in females than males across all outcomes, as seen in the all trials analyses. (G-H) Boundary and non-decision were not different across sexes. Workers: n = 37; slackers: n = 67; males: n = 41; females: n = 63. Bars are mean ± SEM with individual data points laid over the top. *$p<0.05$, **$p<0.01$, ***$p<0.001$.

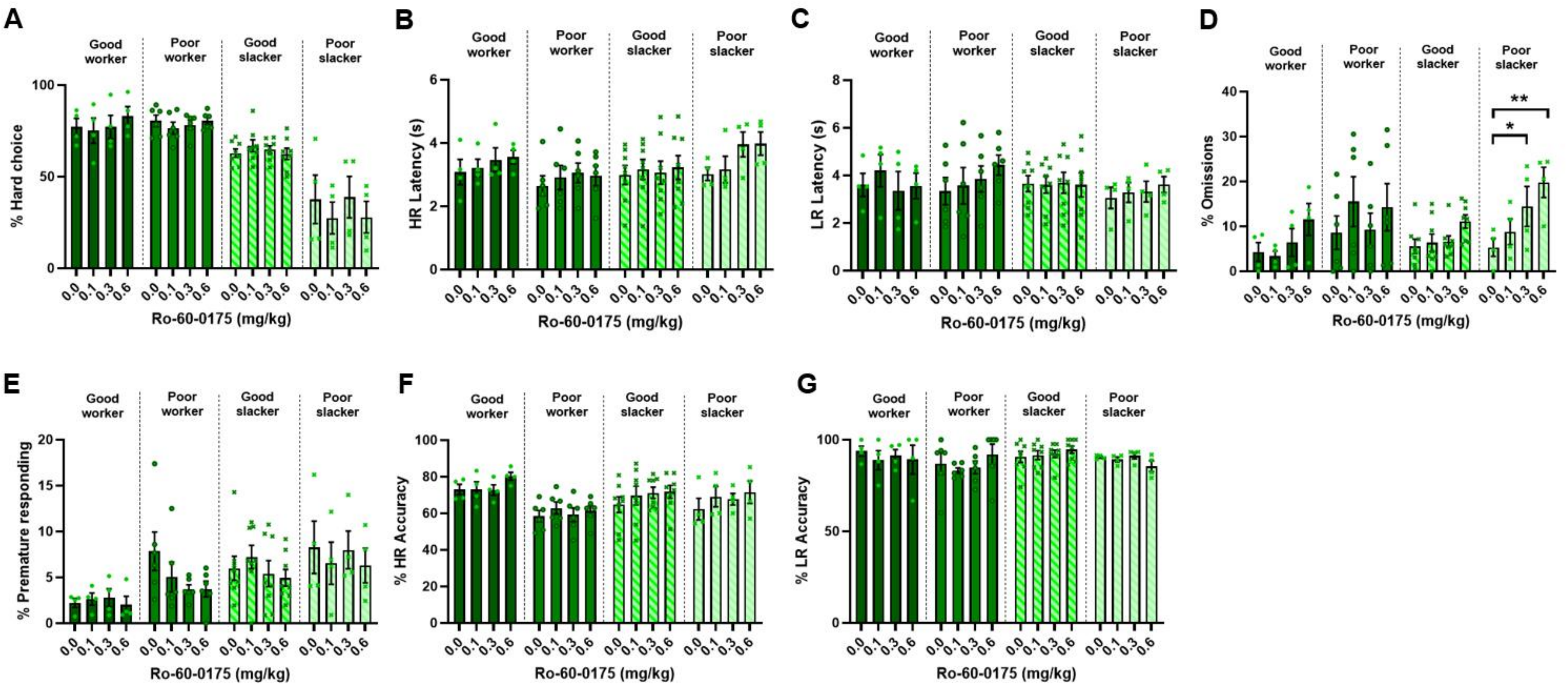


***Supplementary Figure 6 – Reanalysis of rCET behavioral data following administration of Ro-60-0175.***

Behavioral data from the rCET following administration of the 5-$HT_{2C}$ receptor agonist Ro-60-0175 was reanalyzed using four clusters. (A-C) There was no overall effect of Ro-60-0175 on HR choices, or latencies to make choices. (D) When analyzed using clusters, the two highest doses of Ro-60-0175 increased choice omissions in poor slackers only. (E) There was no effect on premature responding. (F) The increase in HR accuracy occurred across all animals. (G) There were no changes in LR accuracy. n=23, bars are mean ± SEM with individual data points laid over the top. *p<0.05, **p<0.01.

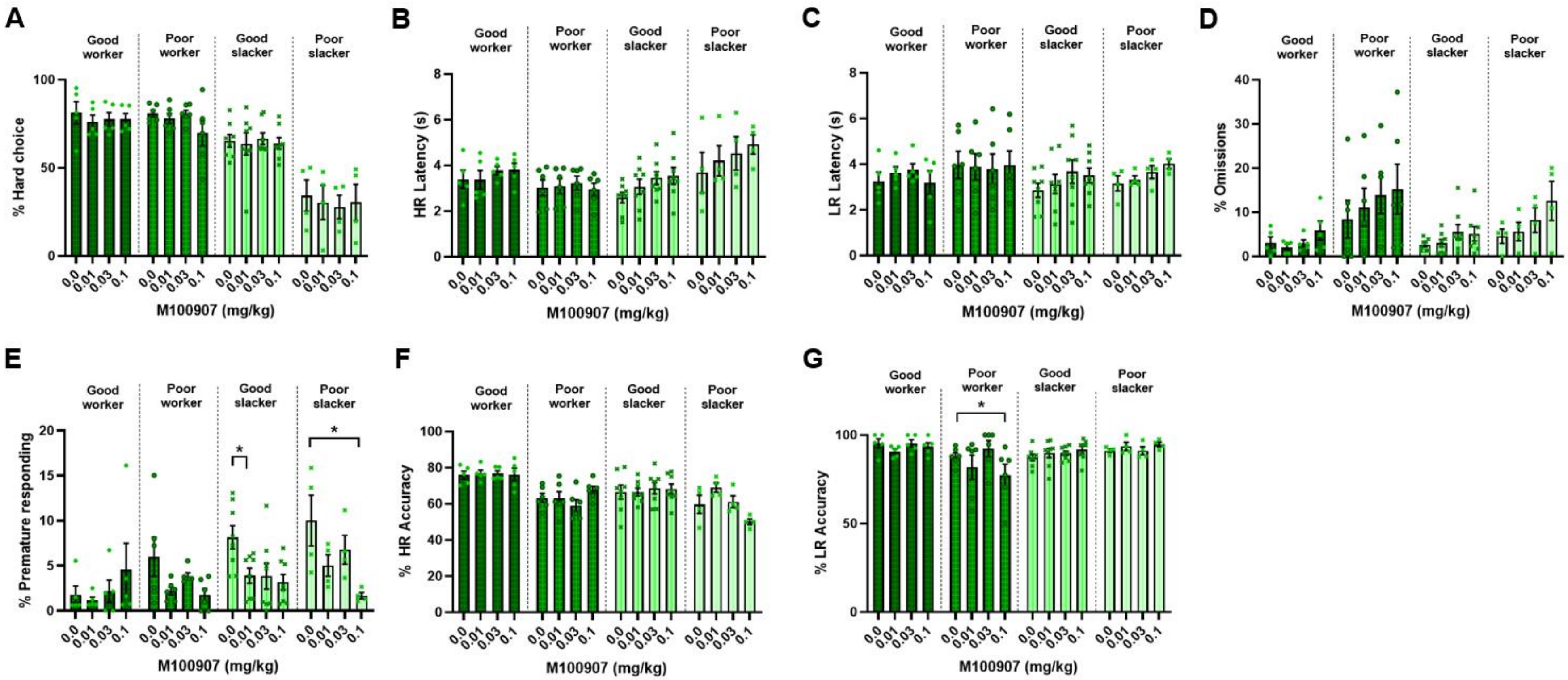


**Supplementary *Figure 7 – Reanalysis of rCET behavioral data following administration of M100907.***

Behavioral data from the rCET following administration of the 5-$HT_{2A}$ receptor antagonist M100907 was reanalyzed using four clusters. (A) There was no effect of M100907 on percentage HR choice. (B-D) The previously found effects of increased choice latencies and increased choice omissions were not specific to cluster, instead the drug had effects across all animals. (E) When analyzed using clusters, the decrease in premature responding was explained by a specific effect in good slackers at the lowest dose, and in poor slackers for the highest dose. (F) There was no effect on HR accuracy. (G) Cluster reanalysis also showed the highest dose of M100907 specifically decreased LR accuracy in poor workers only. n=23, bars are mean ± SEM with individual data points laid over the top. *p<0.05.

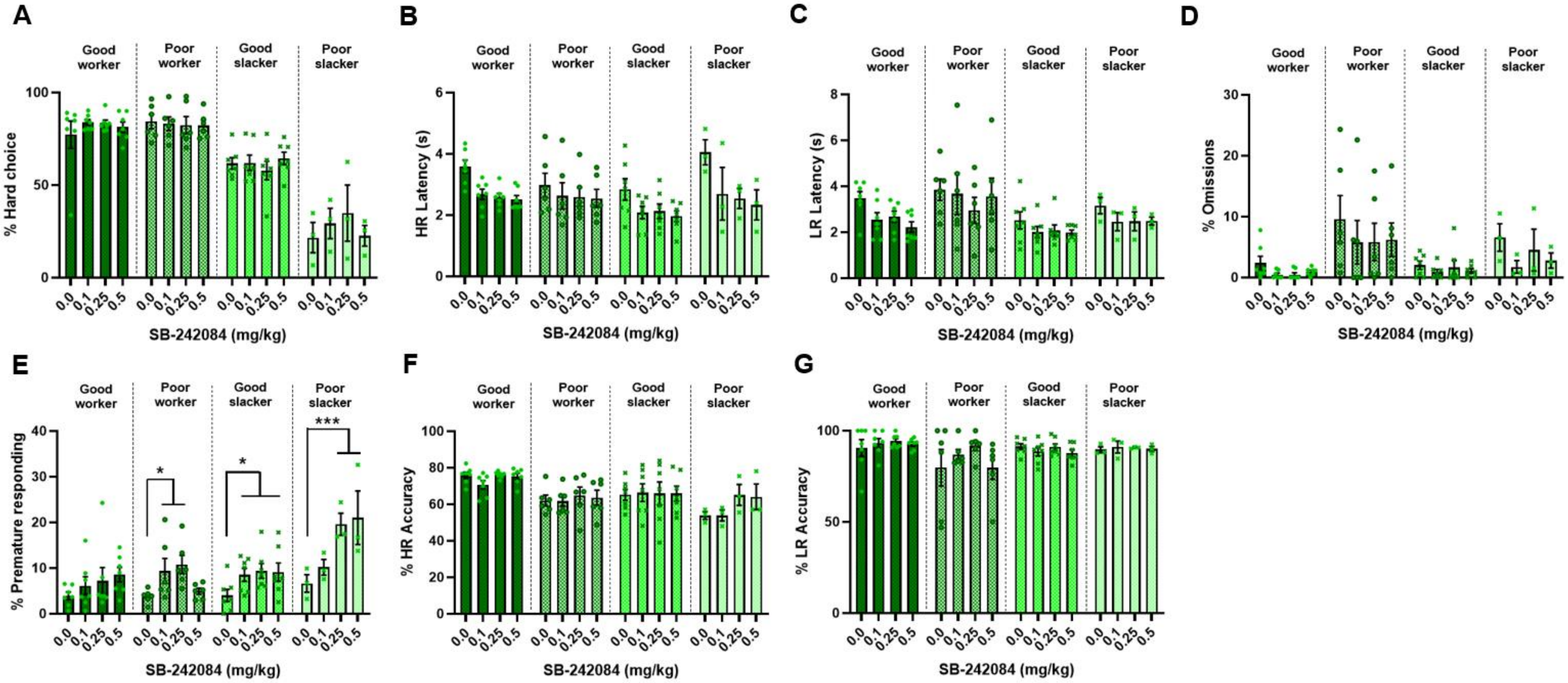


**Supplementary** ***Figure 8 – Reanalysis of rCET behavioral data following administration of SB 242.084.***

Behavioral data from the rCET following administration of the 5-$HT_{2C}$ receptor antagonist SB 242,084 was reanalyzed using four clusters. (A) SB 242,084 did not alter percentage HR choice. (B-C) The previously found effects of decreased choice latencies were not specific to any cluster, instead the drug had effects across all animals. (D) There was no effect on choice omissions. (E) When analyzed using clusters, the increase in premature responding was explained by an effect in all clusters except for good workers. (F-G) HR and LR accuracies were not changed. n=23, bars are mean ± SEM with individual data points laid over the top. *p<0.05, ***p<0.001.

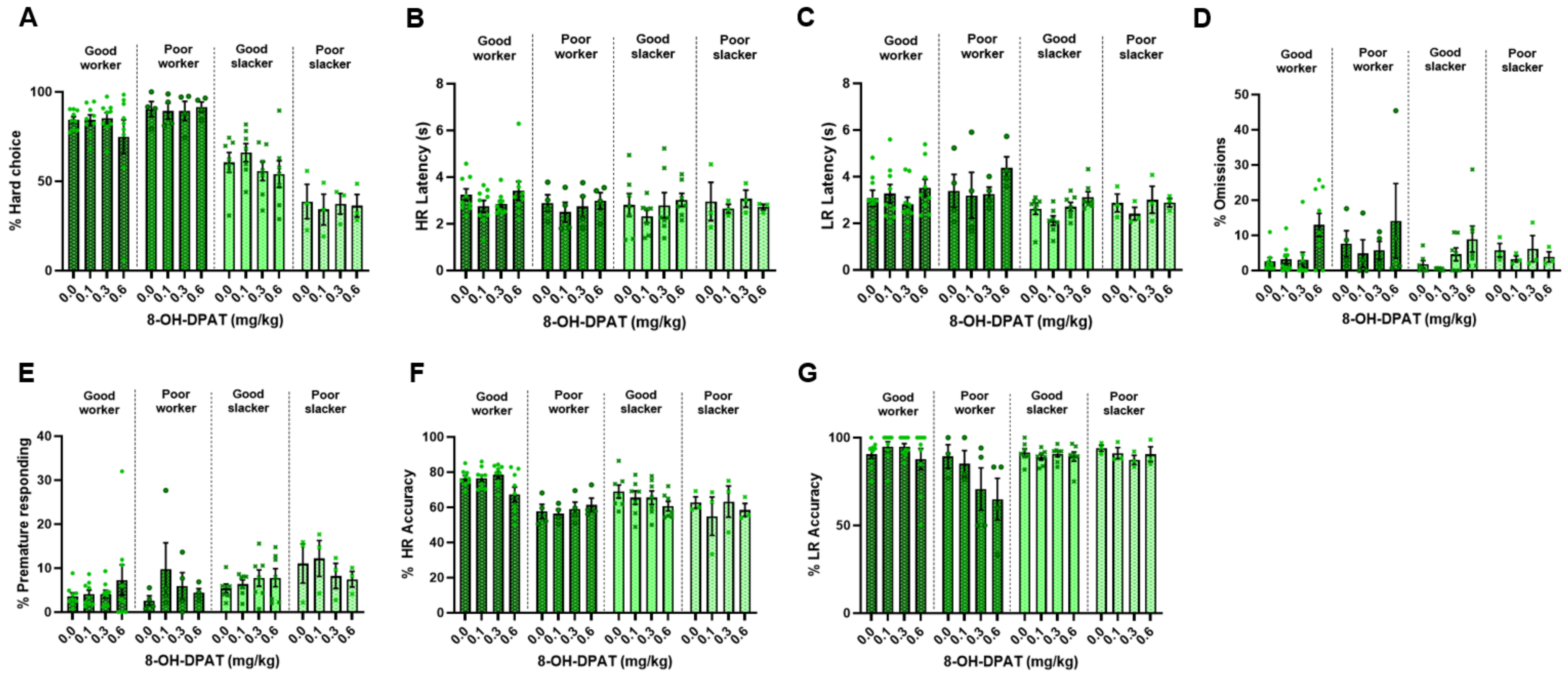


**Supplementary *Figure 9 – Reanalysis of rCET behavioral data following administration of 8-OH-DPAT.***

Behavioral data from the rCET following administration of the 5-$HT_{1A}$ receptor agonist 8-OH-DPAT was reanalyzed using four clusters. (A-G) None of the behavioral changes caused by 8-OH-DPAT were specific to any clusters. Instead, the decrease in LR latency, increase in choice omissions and reduced HR accuracy were explained by effects across all animals. n=23, bars are mean ± SEM with individual data points laid over the top.

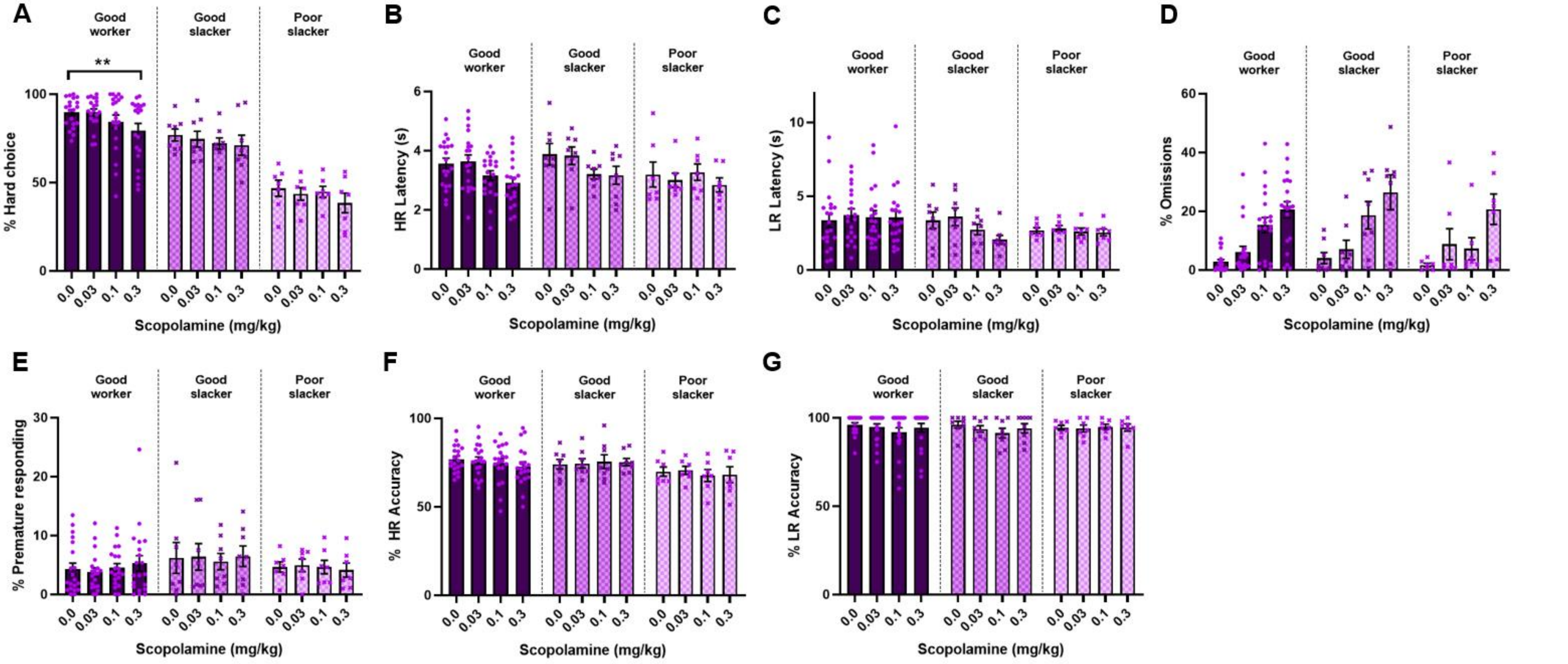


**Supplementary *Figure 10 – Reanalysis of rCET behavioral data following administration of scopolamine.***

Behavioral data from the rCET following administration of the acetylcholine muscarinic receptor antagonist scopolamine was reanalyzed using four clusters. (A) Scopolamine specifically reduced HR choices in good workers. (B) The decrease in HR choice latency was not specific to any cluster. (C) There was no effect on LR choice latency. (D) Scopolamine increased choice omissions across all rats. (E-G) There were no effects on premature responding, HR accuracy or LR accuracy. n=35, bars are mean ± SEM with individual data points laid over the top. **p<0.01.
